# AKT2 modulates astrocytic nicotine responses *in vivo*

**DOI:** 10.1101/2024.05.31.596856

**Authors:** Andrew M. Lombardi, Mina Griffioen, Helen Wong, Ryan Milstead, Curtis Borski, Erin Shiely, Myra E. Bower, Emily Schmitt, Lauren LaPlante, Marissa A. Ehringer, Jerry Stitzel, Charles A. Hoeffer

**Affiliations:** Department of Integrative Physiology, University of Colorado, Boulder, CO 80303; Institute for Behavioral Genetics, University of Colorado, Boulder, CO 80309; Linda Crnic Institute, Anschutz Medical Center, Aurora, CO 80045

**Keywords:** glia, nicotinic receptor, conditioned place preference, Sholl analysis, AKT2, Sholl analysis, conditional knockout, nAChR, Conditioned Place Preference, astroglia, nicotine

## Abstract

A greater understanding of the neurobiology of nicotine is needed to reduce or prevent chronic addiction, ameliorate detrimental nicotine withdrawal effects, and improve cessation rates. Nicotine binds and activates two astrocyte-expressed nicotinic acetylcholine receptors (nAChRs), α4β2 and α7. Protein kinase B-β (Pkb-β or Akt2) expression is restricted to astrocytes in mice and humans and is activated by nicotine. To determine if AKT2 plays a role in astrocytic nicotinic responses, we generated astrocyte-specific Akt2 conditional knockout (cKO) and full Akt2 KO mice. For in/ex vivo studies, we examined mice exposed to chronic nicotine for two weeks in drinking water (200 μg/mL) or following acute nicotine challenge (0.09, 0.2 mg/kg) after 24 hrs. Our in vitro studies used cultured mouse astrocytes to measure nicotine-dependent astrocytic responses. Sholl analysis was used to measure glial fibrillary acidic protein responses in astrocytes. Our data show wild-type (WT) mice exhibit increased astrocyte morphological complexity during acute nicotine exposure, with decreasing complexity during chronic nicotine use, whereas Akt2 cKO mice showed enhanced acute responses and reduced area following chronic exposure. In culture, we found 100 μM nicotine sufficient for morphological changes and blocking α7 or α4β2 nAChRs prevented observed morphologic changes. We performed conditioned place preference (CPP) in Akt2 cKO mice, which revealed reduced nicotine preference in cKO mice compared to controls. Finally, we performed RNASeq comparing nicotine- and LPS-mediated gene expression, identifying robust differences between these two astrocytic stimuli. These findings show the importance of nAChRs and AKT2 signaling in the astrocytic response to nicotine.

**Main Points:**

- Nicotine regulates astrocytes *in vivo*: acute exposure boosts complexity; chronic exposure diminishes it.
- AKT2, expressed in astroglia, modulates morphological changes in response to nicotine and nicotine-dependent conditioned place preference.

## Introduction

Astrocytes are a glial central nervous system (CNS) cell type. Their essential roles include injury responses, synaptic function regulation, and control of ions and neurotransmitters (Barres, 2008; Halassa, Fellin, & Haydon, 2007; Ji, Donnelly, & Nedergaard, 2019; Mahmoud, Gharagozloo, Simard, & Gris, 2019; Perea, Navarrete, & Araque, 2009; Santello, Toni, & Volterra, 2019; Shigetomi, Bowser, Sofroniew, & Khakh, 2008). Under normal conditions, astrocyte state is “naïve” with a diverse stellate morphology. In response to some forms of astrocyte activation, astrocytes undergo shape and functional changes in a process known as reactive astrogliosis. Reactive astrogliosis is part of the CNS response to injury, stress, and disease (M. Pekny & Pekna, 2014; Sofroniew, 2014; A. Verkhratsky, Parpura, Vardjan, & Zorec, 2019). Astrogliosis is characterized by a proinflammatory response that increases the number of glial fibrillary acidic protein (GFAP) positive astrocytes and their morphological complexity (Halassa et al., 2007; Ji et al., 2019; Mahmoud et al., 2019; Milos Pekny & Nilsson, 2005; Perea et al., 2009; Shigetomi et al., 2008; Sofroniew, 2009). In mice, astrogliosis can be induced by lipopolysaccharide (LPS), which triggers an immune reaction through activation of Toll-Like Receptors (TLR) present in astrocytes (Tarassishin, Suh, & Lee, 2014). In recent years, transcriptomic experiments discovered that multiple astrocyte subtypes and responses exist (Batiuk et al., 2020; Hasel, Rose, Sadick, Kim, & Liddelow, 2021; Liddelow et al., 2017; Matusova, Hol, Pekny, Kubista, & Valihrach, 2023; Mizrak et al., 2019; Qian, Qin, Lai, Zhang, & Zhang, 2023; Sun et al., 2024; Taylor et al., 2022). Following the discovery that many forms of “astrogliosis” occur (Anderson, Greenhalgh, Takwale, David, & Vadigepalli, 2017; Batiuk et al., 2020; Ding et al.; Jurga, Paleczna, Kadluczka, & Kuter, 2021; Liddelow et al., 2017; Sun et al., 2024; Vanrobaeys et al., 2023), the term “astrogliosis” now refers to a subset of responses, with most referred to as “activation” (Escartin et al., 2021). Responses to activation are more carefully specified using astrocyte molecular and cellular indicators, such as GFAP, S100β, and EAAT2 (Escartin et al., 2021; Preston et al., 2018; Z. Zhang et al., 2019).

Astrocytes also respond to other forms of neural stimulation, such as the intake of abused substances (Corkrum, Rothwell, Thomas, Kofuji, & Araque, 2019; Periyasamy, Guo, & Buch, 2016), but their role in mediating the neurobiological effects of drugs of abuse is not well understood. Nicotine is one of the most widely used drugs (Jamal et al., 2016) and is known for being associated with the number one preventable cause of morbidity and mortality in the US: tobacco smoking (Jamal et al., 2016). Nicotine can also impact cognition. Withdrawal from nicotine in humans is associated with deficits in memory and learning, which is also seen in mouse models of nicotine dependence (Chellian et al., 2021). Nicotine activates astrocytes in a pattern that is distinct from astrogliosis (Aryal et al., 2021). Astrocytes express two types of nicotinic acetylcholine receptors (nAChRs) alpha7 (α7) and alpha4beta2 (α4β2) (Abbondanza, Urushadze, Alves-Barboza, & Janickova, 2024; Aryal et al., 2021; Gahring, Persiyanov, & Rogers, 2004; Gao, Schneider, Mulloy, & Lee, 2024; Teaktong et al., 2003). These receptors are functionally involved in astrocyte activation during inflammation, neuroprotection, extracellular calcium entry, intracellular calcium release, and hippocampal long-term potentiation (Aryal et al., 2021; Delbro, Westerlund, Bjorklund, & Hansson, 2009; Y. Liu et al., 2012; Ma et al., 2023; Oikawa, Nakamichi, Kambe, Ogura, & Yoneda, 2005; Patel, McIntire, Ryan, Dunah, & Loring, 2017). While upstream astrocyte activators and receptors have been described, the intracellular signaling pathways that regulate morphological changes during astrocyte activation are less well studied. Our study sought to investigate the involvement of astrocyte-specific kinase AKT2 in astrocyte morphological regulation.

AKT (or PKB) proteins are part of a family of serine/threonine kinases with broad cellular roles. AKT signaling controls pathways essential for cell growth, proliferation, metabolism, survival, and memory processes (Cho et al., 2001; Gai, Haan, Scholar, Nicholl, & Yu, 2015; Levenga, Wong, Milstead, LaPlante, & Hoeffer, 2021; J. Zhang et al., 2010). AKT2 is one of three AKT isoforms (AKT1/2/3) found in human and mice brains (Cho et al., 2001; Levenga et al., 2017; Levenga et al., 2021; J. Zhang et al., 2010). Our lab recently demonstrated that AKT2 was the only AKT isoform detected in mouse and human astrocytes *in vivo* (Levenga et al., 2017; Levenga et al., 2021). This finding is not surprising as transcriptomic analysis also shows that AKT2 is the only isoform expressed in astrocytes (Zeisel et al., 2015). Transcriptomic efforts have identified altered brain-related AKT signaling in cancers (Miller et al., 2023; Tamim et al., 2014; J. Zhang et al., 2010), development (Robertson, Coluccio, Jensen, Rydlizky, & Coffman, 2013; D. Zhang, Wang, Zhou, & Xiao, 2017), aging (Kim et al., 2022) and neurodegeneration (Allen et al., 2020; Iacobas, Iacobas, Stout, & Spray, 2020). These findings suggest that AKT2 specifically regulates astrocyte function. A transcriptome-wide association study (TWAS) of smoking phenotypes in humans indicated that AKT2 expression is associated with both cigarettes per day and smoking cessation (Saunders et al., 2022). Additionally, nicotine has been shown to activate AKT signaling (Tsurutani et al., 2005; West et al., 2003) and induce astroglial calcium responses (Araque, Martín, Perea, Arellano, & Buño, 2002; Aryal et al., 2021; Sharma & Vijayaraghavan, 2001). Combined, these data suggest the potential for AKT2 to regulate astroglial responses to nicotine.

Our study sought to determine if nicotine exposure, both *in vivo* and *in vitro,* affected astroglial activation as measured with morpho-functional immunohistochemical analyses, astrocyte activation marker western blot analyses, nicotine-based behavioral tests, and differential gene expression assessments. We also sought to determine if AKT2, the sole AKT isoform expressed in mouse and human astrocytes *in vivo* (Levenga et al., 2017; Levenga et al., 2021), played a role in astrocytic responses to nicotine. Our combined data obtained from hippocampal slice and cell culture preparations indicate that astrocytes respond to nicotine following both acute and chronic exposures. The data also indicate that astrocytic AKT2 activity is required for normal morphological responses to nicotine. We provide evidence that AKT2 is needed for nicotine mediated-conditioned place preference behavior, suggesting that AKT2 function is involved in the manifestation of nicotine-dependent physiological responses. Finally, our transcriptomic data confirms previous observations (Aryal et al., 2021) and supports the possibility that AKT2 may be developed for novel diagnostic and treatment avenues linked to nicotine-use disorders.

## Materials and Methods

### Mice

*Akt2* knockout (KO) mice on a C57BL/6J background were used for this study (Levenga et al., 2017). To generate experimental and control groups, *Akt2^(+/−)^*mice were crossed to generate *Akt2* KO and wild-type (WT) littermates. Mice expressing Cre under the astrocyte *Gfap* promoter (*Gfap*::*Cre*, Jax# 024098) or under the neuroepithelial stem cell *Nes* promoter (*Nes*::*Cre*, Jax# 003771) crossed to floxed *Akt2* mice (Jax# 026475 and (Levenga et al., 2017)) were used for conditional astrocyte-specific or brain-specific KO of *Akt2,* denoted in text as *Gfap-Akt2 cKO* and *Nes-Akt2 cKO,* respectively. The removal of *Akt2* using *Nes*-Cre approaches was confirmed previously (Levenga et al., 2017; Levenga et al., 2021). For this study, AKT2 depletion was confirmed independently using Western blot analysis and immunohistochemical staining and imaging (**Supplemental Figure 1**). *Gfap-Akt2* cKO mice were used for experiments in **Figures 2, 7,** and **8**. *Nes-Akt2* cKO mice were used for experiments in **Figure 5**. *Akt2* KO mice were used for experiments in **Figure 6**. We conducted all experiments with age-matched (4-6 months) and sex-matched littermates. Corresponding N for either biological or technical replicates are described in each figure legend or contained in the figure itself. Mice were kept in cages with sex-matched littermates on a 12:12h light:dark cycle. Lights were turned on at 0700 and off at 1900 with an ambient temperature ranging from 22-24 °C. Food and water were given *ad lib.* Animal care and treatment procedures were approved by the University of Colorado Boulder IACUC and conform to NIH guidelines.

### Primary Astrocyte Culture

Primary mouse astrocyte cultures were generated as previously described (Schildge, Bohrer, Beck, & Schachtrup, 2013). Briefly, cortical tissues were isolated from P0-P4 mouse pups, digested with trypsin, dissociated into single cells, and plated onto T75 flasks coated with Poly-D-Lysine. Cells were grown in astrocyte medium consisting of Dulbecco’s modified eagle medium (DMEM) supplemented with N-2 and 10% fetal bovine serum (FBS). After one week of proliferation, the cultures were shaken at 180 rpm at 37° C for 30 minutes to dissociate adherent microglia, followed by shaking at 240 rpm at 37° C for six hours to remove adherent oligodendrocyte precursor cells. At this point, plates contained enriched astrocytes. After two weeks of growth, astrocytes were re-plated at 0.05×10^6^ cells on 12 mm coverslips for immunostaining. After another 13 days on coverslips, astrocytes were exposed to 100µM nicotine diluted in astrocyte medium, in the presence or absence of 1µM methyl lycaconitine or dihydro-beta-erythroidine, nAChR antagonists, for 24 hours before fixing cells for immunostaining.

### LPS Treatment

Mice were exposed to 5 mg/kg lipopolysaccharide (LPS, Sigma cat# L2630) or vehicle (saline, 0.9% NaCl in water) via intraperitoneal (i.p.) injections. 24 or 72 hours later, mice were sacrificed. The mice were trans-cardially perfused with phosphate-buffered saline (PBS) to remove blood from the brain for imaging and western blotting analysis. One brain hemisphere was post-fixed in 4% paraformaldehyde (PFA) and transferred to 30% sucrose 24-48 hours later. This hemisphere was sliced coronally on a cryostat and stained for GFAP to label astrocytes, Neuronal nuclear protein (NeuN) to label neurons, and Hoechst to label DNA. As described later, GFAP+ astrocytes were used on captured images for morphometric analyses. The other hemisphere was dissected and brain regions, including the hippocampus, were flash frozen for western blot analysis. Brain hemispheres used for biochemistry or imaging were counterbalanced (left or right) for all procedures. The hippocampus was used for staining and western blotting.

### *In vivo* Acute Nicotine Treatment

To test the idea that acute (single exposure) nicotine exposure can modulate astrocyte activation, C57BL/6J (WT) mice were injected intraperitoneally with either vehicle (saline) or nicotine (0.09 or 0.2 mg/kg). 24 hours after treatment, brains were collected for western blotting or immunohistochemistry for morphometric analyses or visualizing astrocyte markers as described below.

### *In vivo* Chronic Nicotine Treatment

To understand the effects of chronic nicotine treatment on astrocytes, mice were given either vehicle (0.2% Saccharin) or nicotine (200µg/µL in 0.2% Saccharin) in their drinking water for two weeks as described in (Mathews & Stitzel, 2019). Mouse weights were collected before all experimental procedures and at the midway point. The average pre-experimental weight was 29.81 g (± 1.81 g), and the average weight at the midway point was 29.69 g (± 1.68 g). The solution was changed four times over the two weeks, and the remaining volume was measured to confirm intake. After the treatment, mice were transcardially perfused, and mouse coronal brain slices were stained for GFAP, S100β, NeuN, and Hoechst. Astrocytes from stained slices were imaged from the CA1 region of the hippocampus using confocal microscopy for GFAP+ or S100β+ astrocytes, and then morphometric analyses were performed as described below.

### Immunostaining

All mouse brains used for immunostaining were fixed by transcardial perfusion using 4% paraformaldehyde (PFA) in PBS and left to fix for 24-48 hours in 4% PFA at 4 °C. Brains were then placed into a 30% sucrose solution for 24 h minimum at 4 °C before embedding in Optimal Cutting Temperature compound (OCT) for sectioning using a Leica cryostat. Coronal 30μm slices were sectioned and stored in cryoprotectant (30% sucrose, 30% ethylene glycol) at −20 °C until use. For immunostaining, slices or coverslips were washed with PBS and blocked for 1 h at 4°C with staining buffer containing 0.05 M Tris pH 7.4, 0.9% NaCl, 0.25% gelatin, 0.5% TritonX-100, and 2% donkey serum. Slices or coverslips were then incubated overnight in the desired primary antibody at 4°C. To detect astrocytes, the GFAP antibody (PhosphoSolutions, 621-GFAP; diluted 1:500) was used as consistent with previous reports (Levenga et al., 2017; Sofroniew, 2009). The NeuN antibody (Novus, NBP1-92693, diluted 1:1000) stained neurons exclusively. This is consistent with previous reports (Gusel’nikova & Korzhevskiy, 2015; Levenga et al., 2017). The AKT2 antibody used (Cell Signaling, 2964, diluted 1:100) stained astrocytes exclusively, consistent with previous reports (Levenga et al., 2017; Levenga et al., 2021). The EAAT2 antibody used (Synaptic Systems 250-203, diluted 1:1000) targets the extracellular domain (Hameed et al., 2023). The S100B antibody used (Invitrogen MA5-12969, diluted 1:100) stained astrocytes exclusively, consistent with previous reports (Jagadeeshaprasad, Govindappa, Nelson, Noble, & Elfar, 2022). After primary incubation overnight at 4°C, slices or coverslips were washed in PBS-T (0.03% TritonX-100) and incubated in the dark at room temperature with fluorescent secondary antibodies (Alexa Fluor 647 conjugated anti-chicken at 1:500; Alexa Fluor 555 conjugated anti-rabbit and anti-mouse at 1:500; Alexa Fluor 488 conjugated anti-mouse IgG1, anti-mouse IgG2A, and anti-rabbit at 1:500; all from Jackson ImmunoResearch) and Hoechst DNA dye at 1:1000 (ThermoFisher). Slices or coverslips were washed again with PBS-T and mounted using Fluoromount-G (ThermoFisher). Z-stack images of the slices and coverslips were taken using the Nikon A1R Laser Scanning Confocal microscope. All laser power and gains were kept the same across slices from the same experiment. Additionally, we imaged the individual channels separately to eliminate any crosstalk. For each slice, 10-20 astrocytes were imaged and selected pseudo-randomly from the CA1 region of the hippocampus. For coverslips, 10-15 astrocytes were pseudo-randomly selected for analysis. Experimenters were blinded during the scoring process for all analyses.

### Sholl Analysis

GFAP-positive astrocytes in the CA1 (stratum radiatum) of the hippocampus at 20x magnification were selected for Sholl analysis using Icy Bioimage Analysis software (ver. 2.3.0.0). 5×5 tiled z-stack maximum projection images were used. The channel for GFAP staining was isolated, and individual astrocytes were selected by zooming in and saving the resulting image. Astrocytes with an apparent nucleus that were not grouped with other astrocytes and had clear GFAP-positive staining were analyzed as randomly as possible by an individual blinded to condition. Images were then binarized to run the Sholl analysis plugin on ImageJ (ver. 1.53f51) on selected astrocytes. In ImageJ, binarization uses a threshold to determine foreground vs. background. Foreground pixels are colored black, and background pixels are colored white. The threshold is determined by finding the number in the threshold editing window that preserves the most contiguous astrocyte processes from a subset of images from each condition. This number is then used to binarize all images using batch settings. Any errant pixels that would distort Sholl analysis are colored white, resulting in their removal from analysis. Astrocyte images are then run through the Sholl analysis ImageJ plugin using the following parameters: Starting Radius: 0μm, Step Size: 10μm, and End Radius: 125μm. Batch settings reduce the time it takes to run the analysis, and each image is run through the plugin individually. Data are then graphed as the number of intersections versus radius from the astrocyte soma. Some confocal images captured early in the study used a 40X objective. When this equipment later became unavailable, images were captured using a 20X objective. To confirm imaging using either configuration was indistinguishable, we compared these different sets, normalizing them to % of saline controls. No differences were found (**Supplemental Figure 2A, B**).

### Area Analysis

Astrocytes (from either coverslips or brain slices) were binarized as described during preparation for Sholl analysis. Astrocytes are then run through the area measure feature in ImageJ. The area of the black pixels in the image is calculated and data are given as pixel units squared. Area was calculated from GFAP staining, except for the addition of S100β staining on astrocytes exposed to chronic nicotine. Areas were then represented as % of vehicle controls. 3 slices were used per animal, with 10-20 astrocytes per slice. For *in vitro* analysis, 4 coverslips were used per condition with 10-15 astrocytes per coverslip. Analysis was done by an individual blinded to condition.

### Intensity Analysis

For sum intensity measurements, Z-stacks imaged from mouse hippocampal slices stained at the same time, were maximum projected using ICY Imaging Software. After maximum projection, sum intensity was measured in the CA1 region of the hippocampus using the region of interest (ROI) tool. ROIs of identical areas were used on all slices. 2-3 slices were used from each animal, 3 animals per condition. Analysis was done by an individual blinded to condition.

### Hippocampal Slice preparation

To examine AKT2 activation, acute 400 µm transverse hippocampal slices were prepared as previously described (Levenga et al., 2013; Wong et al., 2015). Hippocampal slices were derived from WT male mice 8-12 weeks of age. Briefly, slices were maintained in an interface chamber at 32°C infused with oxygenated artificial cerebral spinal fluid (ACSF) containing (in mM): 125 NaCl, 2.5 KCl, 1.25 NaH2PO4, 25 NaHCO3, 25 D-glucose, 2 CaCl2 and 1 MgCl2. Slices recovered in the chamber at least 60 min before nicotine or vehicle application. Next, vehicle (Saline) or nicotine (100 μM) was applied to slices for 30 minutes. At the end of the treatment period, slices were flash frozen on dry ice, and lysates prepared as described below for Western blotting.

### Western blotting and total protein measurements

Protein lysates were obtained using procedures as previously described (Levenga et al., 2017). Mouse hippocampi were homogenized via sonication in lysis buffer containing: 10mM HEPES pH 7.4, 150 mM NaCl, 50mM NaF, 1mM EDTA, 1mM EGTA, and 10mM Na_4_P_2_O_7_, including 1x protease inhibitor cocktail (SIGMA P8340-5ML), and phosphatase inhibitor cocktails II and III (SIGMA P5726-5ML and P0044-5ML). 20µg samples were made in Laemmli sample buffer and separated by size on 4-12% Bis-Tris gradient gels. Proteins were then transferred to PVDF membranes. These blots were blocked with 0.2% I-Block (Tropix T2015) dissolved in Tris-buffered saline with 0.1% Tween-20 (Acros Organics 23336-0010). Blots were incubated with primary antibodies diluted in I-Block for 24 hours at 4°C. The GFAP antibody used (PhosphoSolutions, 621-GFAP; diluted 1:20,000) recognized only the expected band at 50 kilodaltons (kDa) on western blots of mouse brains. The total AKT2 antibody used for western blot analysis (LS Bio, LS-C156232, diluted 1:2500) recognized the expected band at 60 kDa for mouse brains, as previously shown (Levenga et al., 2017). This antibody has a non-specific band below the confirmed AKT2 protein band. We verified the antibody by blotting *Akt2* KO tissue, resulting in no antibody banding at the expected molecular weight, so the non-specific band was excluded from analysis. The phospho-AKT2 Ser474 (pAKT2) antibody (Cell Signaling, 8599; diluted 1:750) recognized only the expected band at 60 kDa on western blots of mouse brains, consistent with previous reports (Levenga et al., 2017). The GAPDH antibody (Cell Signaling, 5174, diluted 1:1000) recognized only the expected band at 37 kDa on western blots of mouse brains, consistent with previous reports (Levenga et al., 2017; Levenga et al., 2021; Shi et al., 2023). GAPDH was used as a loading control. The Occludin antibody (Invitrogen OC-3F10, 1:1000) detected only the expected band at 53 kDa in westerns of mouse brains, as consistent with previous reports (Sole et al., 2023). The ALDH1L1 antibody (Novus NBP2-50033, diluted 1:5000) recognized only the expected band at 100 kDa on western blots of mouse brains and stained astrocytes exclusively, consistent with previous reports (Levenga et al., 2021). The S100B antibody (Cell Signaling 9550, diluted 1:500; (Huang et al., 2023)) detected only the expected band at 9 kDa in western blots of mouse hippocampus, as consistent with previous reports (Ebert et al., 2021). The Vimentin antibody for WB (Abcam, ab92547, diluted 1:1000) detected only the expected band at 55 kDa in westerns of mouse brains, consistent with previous reports (Bakshi et al., 2016). To detect primary antibodies, blots were incubated with secondary HRP-conjugated goat anti-mouse (Promega W402B), anti-rabbit (Promega W4011) or anti-chicken (Abcam ab97135) antibody (at 1:5000 in I-Block or same concentration as primary if more dilute). Signal was detected using chemiluminescence and protein levels quantified as previously described (Levenga et al., 2017). Loading controls and total protein were used to normalize the signal.

Per the manufacturer’s guidelines, total protein measurements were performed using Azure TotalStain Q-PVDF (Cat # AC2225). In brief, after gel-separated protein was transferred to PVDF membranes, blots were washed with ddiH2O for 5 minutes with gentle agitation on an orbital shaker at room temperature. Water was decanted, and the blot was incubated in PVDF Staining Solution (Azure S1071) for 5 minutes with gentle agitation at room temp. The staining solution was decanted, and the blot was washed 3 times in Washing Solution (Azure S1072) with gentle agitation at room temp for 6 minutes each. The blot was then placed into the imaging folder (Azure S1092), and total protein was visualized with an Azure Biosystems 600 Imaging Systems using the Cy3 laser channel. Blots were then rinsed with ddiH2O and blocked with 0.2% I-Block w/ 0.1% Tween-20 at room temperature for 1 hour for subsequent primary antibody incubation.

### Conditioned Place Preference

Conditioned place preference (CPP) procedures were adapted from (MG. Kutlu, LA. Ortega, & TJ. Gould, 2015). Briefly, the CPP apparatus was set up using three connected 20 cm x 40 cm plexiglass chambers with removable dividers connecting each. Each chamber is differentiated by different wall colors and flooring materials (wire mesh of differing composition). The testing light conditions were 585 lux, with a background sound of 62.5 dB, which were maintained during all phases of the CPP procedure. For CPP testing, mice were placed in the center chamber and allowed to explore the CPP apparatus for 15 minutes. Time spent in each chamber was measured with the Ethovision XT17 video tracking system (Noldus) to determine baseline chamber preference for preconditioning. Starting the next day, mice received two daily subcutaneous (SQ) injections 5 h apart for CPP training over 3 consecutive days. Mice in the nicotine group received a nicotine injection at 0.35 mg/kg and saline injection 5 h apart daily. After the nicotine injection, mice were placed for 15 minutes in the CPP chamber that they did not prefer on the preconditioning day while saline injections were given to the nicotine test group in the preferred chamber. This nicotine treatment group is noted as ‘Nicotine/Saline’. The order in which nicotine and saline were given for the nicotine group mice was randomized each day. Control mice received two saline injections 5 h apart and were randomly placed in either chamber for each training day, regardless of whether chamber was preferred or non-preferred. This control treatment group is described as ‘Saline/Saline’. After 3 days of conditioning, mice were allowed full access to both chambers, and the time spent in each chamber was analyzed with Ethovision. Entry to the chamber was only scored once the whole body, including hindlimbs, was in the zone. The starting chamber was randomized before testing. There were four groups total: WT Nicotine/Saline, WT Saline/Saline, KO Nicotine/Saline, and KO Saline/Saline. Preference score was determined by subtracting the time spent in the unpreferred chamber on preconditioning day from the time spent in the unpreferred chamber on testing day 2. Time spent in the unpreferred chamber and distance traveled during testing were measured to confirm validity of results. Data are shown as the distance moved or time spent in the unpreferred chamber by each individual nicotine-treated animal subtracted by the average saline score from each genotype. No sex-dependent differences were seen; therefore, we combined both male and female data.

### Transcriptomics

To identify differences in gene expression between LPS and nicotine exposure, we performed RNA sequencing. WT male mice 2-6 months of age were treated with 5mg/kg of Lipopolysaccharide (LPS) (N=4) or 0.9% Saline solution (N=4) through intraperitoneal injection, prefrontal cortex (PFC) was dissected after 24 hours, and LPS samples were compared to saline for gene expression analysis. For nicotine studies, mice 2-6 months of age were given 0.2 mg/kg nicotine (N=6) or 0.9% saline (N=6), the hippocampus was dissected after 24 hours, and Nicotine samples were compared to saline for gene expression analysis. Standard techniques were used for RNA isolation, sequencing, and gene expression analyses. Briefly, tissue was homogenized with a glass Dounce homogenizer in Trizol reagent, followed by RNA extraction following the manufacturer’s instructions. Poly(A)-selected libraries were created using the Universal Plus™ mRNA-Seq library preparation kit with NuQuant. Libraries were sequenced on Illumina NovaSeqX to produce >40M 150bp paired-end FASTQ reads. Reads were trimmed and filtered for quality using fastp (S. Chen, Zhou, & Chen, 2018), then aligned to the mm10 *Mus Musculus* genome using STAR (Dobin et al., 2013). Raw read/gene counts for alignment files were obtained using featureCounts (Liao, Smyth, & Shi, 2014), and differential gene expression was analyzed using DESeq2 software (Love, Huber, & Anders, 2014) with a significance cut-off of BH-corrected adjusted p<0.1. Gene ontology over-representation test was performed using the ClusterProfiler package (G. Yu, L. Wang, Y. Han, & Q. Y. He, 2012) to examine gene expression profiles with a significance cut-off of p-value<0.1. Transcriptomic results were uploaded to the Gene Expression Omnibus (GEO) database and are available in **Supplementary Tables 1** and **2).**

### Statistics

GraphPad (ver 9.1.2) software was used to perform statistical analyses. All data are shown as mean ± standard error of the mean (SEM). Differences between groups were determined with either student’s t-test or N-way ANOVA. Outliers were identified and removed using the ROUT method. If ANOVA showed statistical significance, it was followed by Tukey’s post hoc testing. All statistic tests were two-tailed with an alpha of p<0.05. When data did not have equal variances, Welch’s t-test was used. For Sholl analysis statistics, a 2-way ANOVA mixed model was used with the following variables: Sholl Radius, Group, and Interaction. For area analysis, a 2-way ANOVA mixed model was used with the following variables: Genotype, Treatment, and Interaction. For CPP, a t-test was used to compare WT nicotine preference to *Gfap-Akt2* cKO nicotine preference relative to saline controls. Exact p-values are reported in the text, except when p<.0001, then that number is used instead. In Figures, simple p-values are reported as symbols defined in corresponding figure legends. Comprehensive summary statistics are included in Table 1.

## Results

### Sholl and area analysis can detect morphological changes consistent with astrogliosis

We initially tested our ability to detect astroglial morphological changes in the mouse brain induced by experimental manipulation, so we exposed mice to LPS *in vivo* (5 mg/kg) to induce astrogliosis. Using morphometric analyses, we were able to detect significantly increased size and morphological complexity of astrocytes in the hippocampus of LPS-treated mice compared with vehicle-treated (saline) control mice. The average area of astrocytes defined by GFAP was larger in LPS-treated mice than in control mice at each time point assessed (**Figure 1A, B**; main effect of LPS treatment: p<0.0001; post-hoc testing, significant differences between Saline and LPS at 24h and 72h: p<0.0001 for both time points). Similarly, astrocyte morphological complexity measured via Sholl analysis showed a significant increase in astrocyte branching at each time point (**Figure 1A, B**, main effect of LPS treatment: p<.0001; post-hoc testing, significant differences between Saline and LPS at 24h and 72h: p<0.0001 for both time points). See Table 1 for specific statistics. Western blot analysis of GFAP expression using hippocampal lysates obtained from these mice did not reveal any changes in GFAP levels between LPS treatments and saline (**Figure 1C**). This was not necessarily unexpected as previous studies have reported no difference in GFAP levels with LPS (Norden, Trojanowski, Villanueva, Navarro, & Godbout, 2016; Tarassishin et al., 2014). However, we did detect LPS-mediated increases in *Gfap* mRNA (**Supplemental Figure 3A**) as well as increased gene expression of signature astrogliosis markers (**Supplemental Figure 3B**) using transcriptomic methods. Overall, we observed a total of 4,257 differentially expressed genes, with 2,330 genes upregulated and 1,927 genes downregulated by LPS treatment (**Supplementary Table 1**). This gene expression data combined with our area and Sholl morphometric analysis parameters confirm our ability to detect LPS-induced morphological alterations and activation of astrocytes following *in vivo* stimulation.

**Figure 1:**
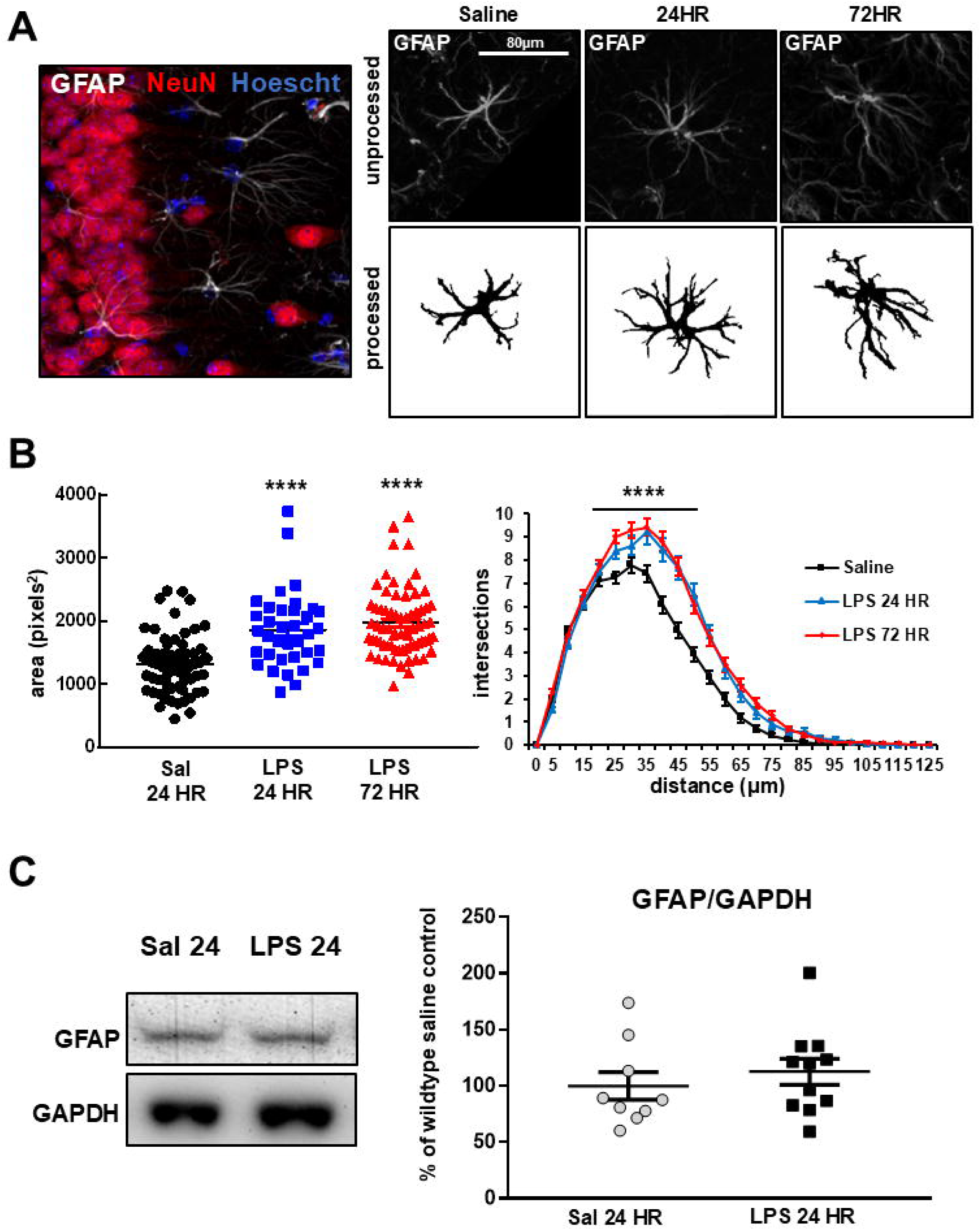
Confirmation of Sholl analysis using LPS-induced inflammatory insult. Hippocampal astrocytes in WT mice were assessed for morphological changes 24 h and 72 h post-LPS treatment (i.p.) using area and Sholl analyses. (**A**) Representative images used for analysis of WT astrocytes in hippocampal area CA1 after LPS treatment. *Left:* Confocal images showing neurons (NeuN, red), astrocytes (GFAP, white) and nuclei (Hoechst, blue). *Top Right:* Unprocessed representative images of hippocampal astrocytes (GFAP, white). *Bottom Right:* Processed (Binarized) images of astrocytes. **(B)** LPS induces GFAP-dependent increases in area and Sholl complexity. **** p<0.0001 for all comparisons except 24-hour LPS vs. 72-hour LPS. ANOVA, Tukey’s *post hoc*. **(C)** GFAP western blot staining of hippocampal lysate. No significant difference in total GFAP levels was detected between LPS-treated and control groups. p >.05 for all comparisons. Glyceraldehyde-3-phosphate dehydrogenase (GAPDH) is used as a loading control. Representative western blot images used for the generation of quantitation graphs. Imaging data is representative of 3 stained slices per mouse and 10 astrocytes/mouse. Biological N=3 for each group. For Westerns, N=3 for each group. Groups were alternated by sex, either 2M, 1F, or 2F, 1M. ANOVA. See Table 1 for exact statistics.

### Conditional knockout (cKO) of *Akt2* in astrocytes leads to increased morphological response to LPS treatment

Because we found in earlier studies that AKT2 protein was restricted to astrocytes (Levenga et al., 2021; Wong et al., 2020), we next sought to determine if AKT2 activity regulated astrocytic responses to LPS using an astrocyte-specific *Akt2* cKO (*Gfap-Akt2* cKO) mouse. AKT2 removal and astrocyte specificity were confirmed using immunohistology and western blotting of *Akt2* cKO brain tissue (**Figure 2A**; p<0.0001). We then assessed whether astrocytes of *Akt2* cKO mice respond differently than in WT mice during LPS-induced astrogliosis. At 24h post-LPS treatment, *Akt2* cKO astrocytes showed significantly increased size and morphologic complexity when compared with WT LPS or control groups of either genotype. Additionally, saline-treated *Akt2* cKO astrocytes displayed a small but significantly increased degree of morphological complexity compared with saline-treated WT controls (**Figure 2B, C**; interaction effect for treatment x genotype on Sholl distance: p<0.0001; post hoc testing, WT Saline vs. WT LPS: p=0.0489, WT Saline vs. *Akt2* cKO LPS: p<0.0001, WT LPS vs. *Akt2* cKO LPS: p<0.0001, WT Saline vs. *Akt2* cKO Saline: p=0.0104, *Akt2* cKO Saline vs. *Akt2* cKO LPS: p=0.0001). Similarly, the area of astrocytes defined by GFAP staining in both *Akt2* cKO and WT mice increased with LPS treatment, but there was no difference in GFAP-dependent area between genotypes in saline treatment groups, contrasting the Sholl results (**Figure 2B, D**; main effect of treatment: p<0.0001, main effect of genotype, p=0.0254; post hoc testing, WT Saline vs. WT LPS: p=0.050, *Akt2* cKO Saline vs. *Akt2* cKO LPS: p=0.0016, WT Saline vs. *Akt2* cKO LPS: p<0.0001, WT Saline vs. *Akt2* cKO Saline: p=0.1539). See Table 1 for specific statistics. These results provide evidence that AKT2 activity is involved in morphological responses to LPS-induced astrogliosis, and its absence may enhance expression of LPS-dependent astroglial morphological changes.

**Figure 2:**
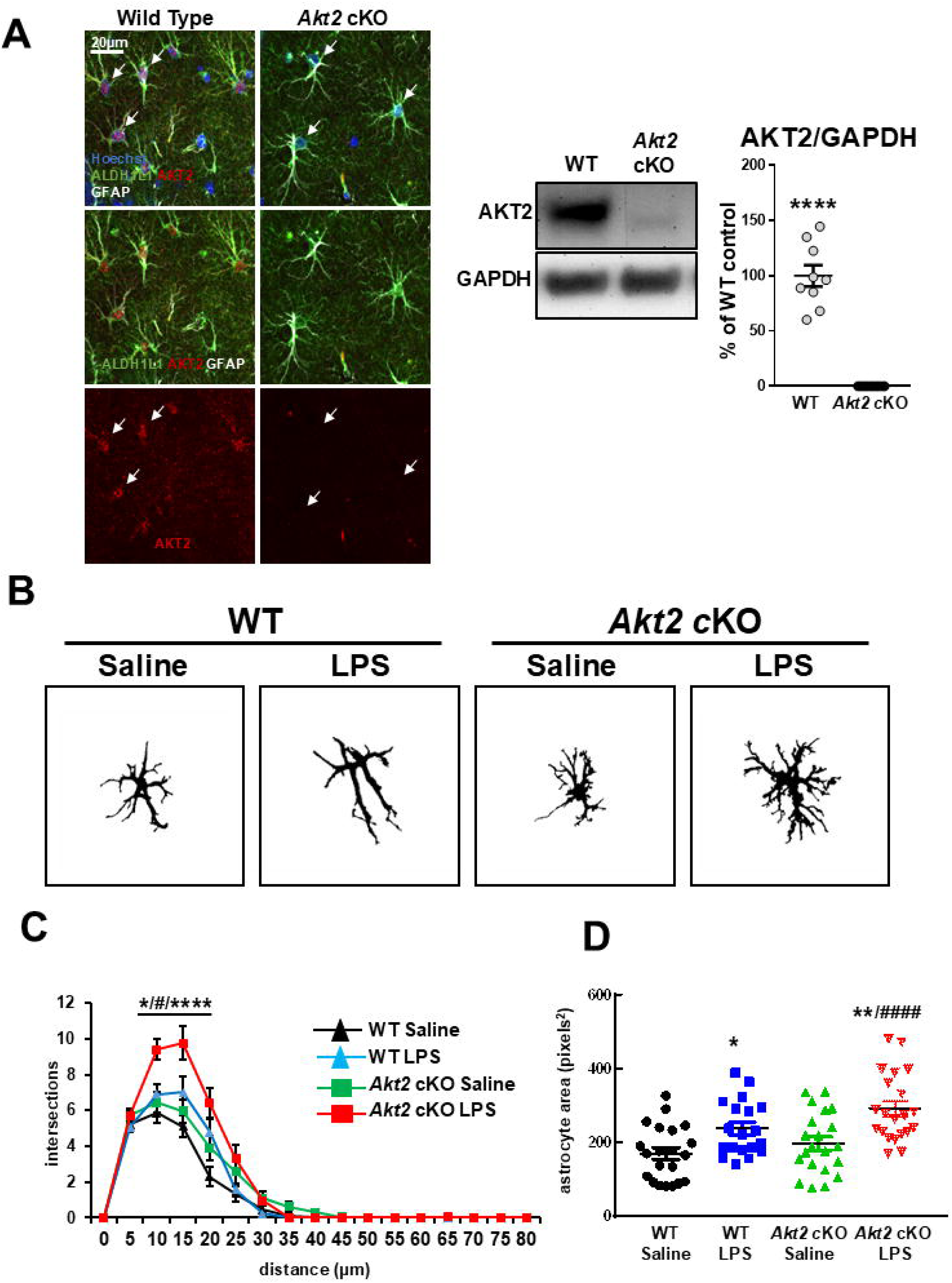
Conditional removal of *Akt2* from astrocytes enhances astrocyte activation in response to LPS treatment. **(A)** WT and *GFAP-Akt2* cKO (*Akt2* cKO) mice were exposed to LPS and sacrificed 24 hours later. Immunohistochemical staining confirms the removal of AKT2 from astrocytes in *Akt2* cKO mice (left). Western blotting to confirm expected AKT2 levels (right), **** p<.0001. **(B)** Representative binarized images of astrocytes in the hippocampus of WT and *Akt2* cKO mice treated with LPS or saline. These images were used for morphometric assessments of astrocytes using **(C)** Sholl analysis and **(D)** GFAP-dependent area analysis. For Sholl analysis; WT Saline vs WT LPS: * p<0.05; *Akt2* cKO Saline vs. *Akt2* cKO LPS, WT Saline vs. *Akt2* cKO LPS, WT Saline vs. *Akt2* cKO: **** p<0.0001; WT Saline vs. *Akt2* cKO Saline: # p<0.05. For GFAP-dependent area analysis; WT Saline vs WT LPS: * p=0.050; *Akt2* cKO Saline vs. *Akt2* cKO LPS: ** p<0.01; WT Saline vs. *Akt2* cKO LPS: #### p<0.0001. Imaging data representative of 3 stained slices per mouse and 10 astrocytes/mouse. Biological N=3 for each group. Groups were alternated by sex, either 2M, 1F, or 2F, 1M. For Westerns, N=9 for each genotype and 5M and 4F for each group. ANOVA, Tukey’s post hoc testing. See Table 1 for exact statistics.

### Astrocytes display dose-dependent increases in morphological complexity after acute nicotine exposure

Having established our ability to detect astrocyte morphometric responses to LPS, we next assessed the effect of *in vivo* nicotine exposure on astrocytes. We acutely treated WT mice with a single dose of nicotine (0.09 or 0.2 mg/kg) via i.p. injection. At 24 hours post-nicotine exposure, we found that astrocytes became significantly larger and morphologically more complex (**Figure 3A, B;** main effect of nicotine treatment, p<0.0001, post hoc group comparisons; p<0.0001 for all group comparisons), consistent with astrocyte activation (Aryal et al., 2021; Escartin et al., 2021). Using area analysis, we also observed significant increases in GFAP-defined astrocyte area following nicotine treatment *in vivo,* which was dose-dependent (**Figure 3C**, main effect of treatment, p<0.0001; post hoc testing, Saline vs. 0.09 mg/kg Nicotine: p=0.0020, Saline vs 0.2 mg/kg Nicotine: p<0.0001, 0.09 mg/kg nicotine vs 0.2 mg/kg nicotine p=0.0395). No difference in total numbers of astrocytes/area was detected for any treatment group (**Figure 3D**), indicating that Sholl and area measurements were not confounded by unequal numbers of imaged astrocytes.

**Figure 3:**
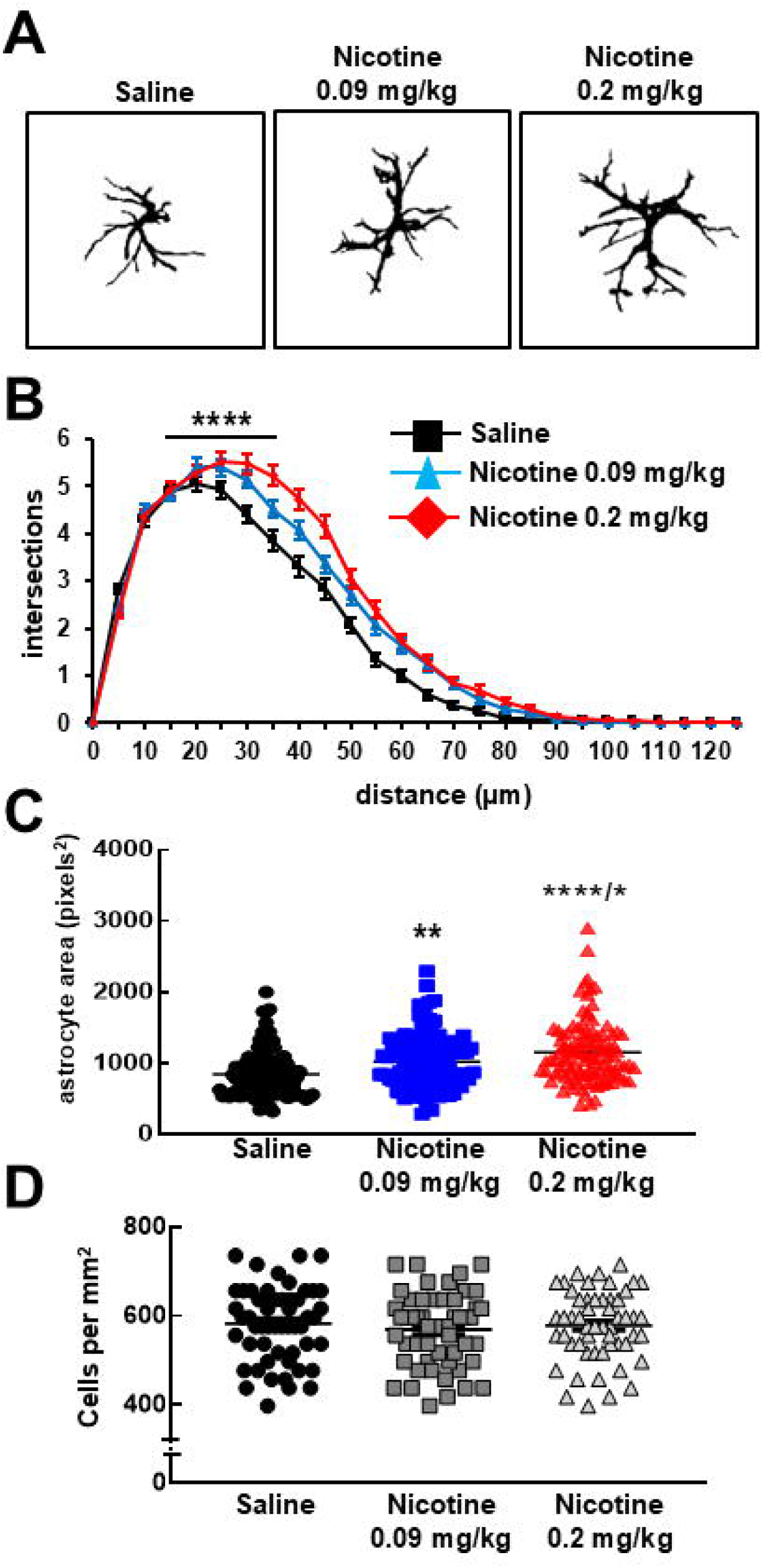
*In vivo* acute nicotine exposure promotes astrocyte activation. Sholl analysis of hippocampal astrocytes following acute nicotine treatment (0.09 and 0.2 mg/kg *in vivo*) measured 24 hours after exposure compared to saline injection. **(A)** Representative image of binarized astrocytes for each treatment condition. **(B)** Sholl quantification of WT responses to acute nicotine treatment show a dose-dependent increase in astrocyte morphological complexity 24h after nicotine exposure. Post hoc comparisons for all groups: **** p<0.0001. **(C)** GFAP-based area analysis shows dose-dependent increases in astrocyte size 24h post-nicotine exposure in WT mice. Saline vs. 0.09mg/kg: ** p<0.01; Saline vs. 0.2 mg/kg: **** p<0.0001, 0.09 mg/kg vs. 0.2 mg/kg: * p<0.05. **(D)** Cell counts for each experimental group. No differences between groups were detected. For sholl analysis, p values <.001 for all group comparisons. Imaging data representative of 3 stained slices per mouse and 15 astrocytes/mouse. Biological N=3 for each group. All mice were male. See Table 1 for exact statistics.

### Acute nicotine treatment regulates some markers of astrocyte activation and activates AKT2

After determining that acute *in vivo* nicotine exposure promoted morphological alterations in astrocyte complexity and area, we next examined the expression of astrocyte activation markers. Many astrocyte activation markers have been identified and described (Escartin et al., 2021). GFAP and excitatory amino acid transporter 2 (EAAT2), also known as solute carrier family 1 member 2 (SLC1A2) or glutamate transporter 1 (GLT-1), have been linked to nicotine. A prior study examining nicotine effects on astrocytes found increased GFAP levels using quantitative imaging (Aryal et al., 2021), while EAAT2’s relationship to nicotine use has been established in preclinical models and in humans (Knackstedt et al., 2009; McClure, Gipson, Malcolm, Kalivas, & Gray, 2014). We did not detect significant differences in GFAP protein levels using Western blot analysis of hippocampal lysates from animals that were acutely treated with nicotine (**Figure 4A)**. *Gfap* mRNA expression also showed no significant differences with acute nicotine treatment (**Supplemental Figure 3C**). Based on findings from Aryal et al. 2021, we next assessed GFAP levels using immunohistochemical staining of the hippocampus. Using this approach, a small but significant increase in GFAP levels in the nicotine treatment group was detected 24 hours after treatment compared to controls (p=0.0470, **Figure 4B**). With EAAT2, we also did not detect significant differences in protein levels between treatment groups by Western blot analysis (**Supplemental Figure 4A)**; however, staining and image quantification of the hippocampus revealed that nicotine-treated mice displayed ∼40% reduction in EAAT2 staining intensity compared with saline-treated controls (p<0.0001, **Figure 4C**). We also examined total levels of several other known markers of astrogliosis and astrocyte activation (Escartin et al., 2021). Western blotting analysis of hippocampal lysate from nicotine- and saline-treated mice revealed no significant protein expression differences in several astrocyte activation markers (**Supplemental Figure 4**). Additionally, unlike LPS, acute nicotine exposure did not significantly alter the gene expression of several astrogliosis markers (**Supplemental Figure 3D**). These data confirm that acute nicotine exposure alters the immunohistochemically determined expression of some astrocyte activation markers, but the nicotine-activation marker profile is different from astrogliosis (Aryal et al., 2021; Escartin et al., 2021).

**Figure 4:**
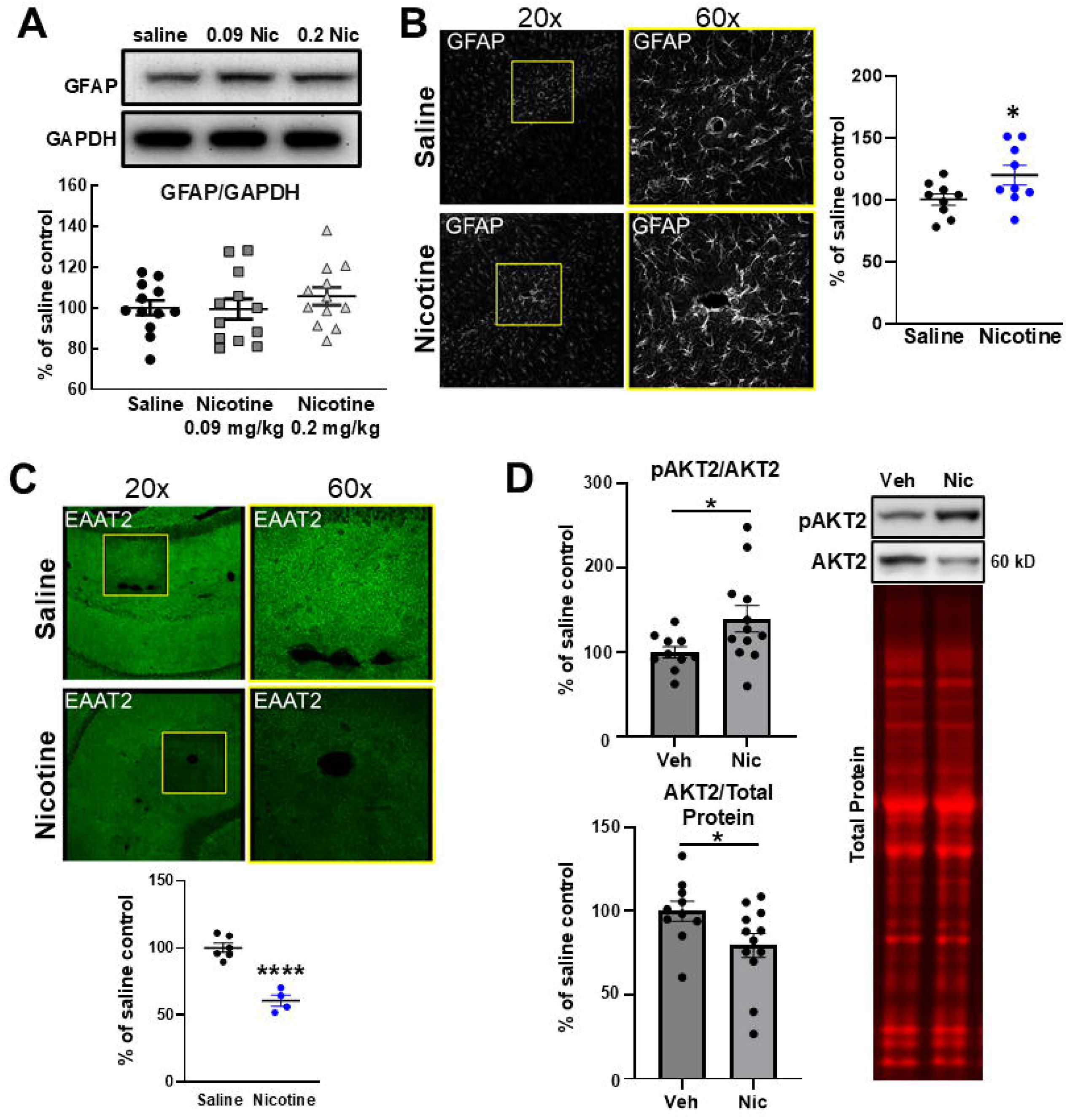
Astrocytic marker expression is regulated by *in vivo* nicotine exposure. Representative staining images of astrocyte marker expression from mouse hippocampus (area CA1) following i.p. injection after 24 hours of 0.2 mg/kg nicotine. AKT2 activation was measured after 100µM nicotine treatment to *ex vivo* hippocampal slices. **(A)** Western blot analysis of mouse hippocampal lysate revealed no significant change in GFAP levels following acute nicotine treatment, although trends were seen (see Table 1). Glyceraldehyde-3-phosphate dehydrogenase (GAPDH) was used for loading controls. ANOVA. **(B)** GFAP (white) staining is increased following acute nicotine exposure. Quantification of staining shows significantly increased staining intensity of GFAP, * p<0.05. T-test. **(C)** EAAT2 (green) staining is reduced following acute nicotine exposure. Quantification of staining shows significantly reduced EAAT2 staining intensity **** p<0.0001. T-test. **(D)** AKT2 is activated in *ex vivo* WT hippocampal slices after 30 min following 100μM nicotine treatment. Phospho-AKT2/Total AKT2 ratios (pAKT2, Serine 474) were higher in nicotine-treated hippocampal slices than in controls. * p<.05. Total AKT2 levels were also reduced 30 min post nicotine treatment. * p<.05. Representative blot next to quantification. No differences in total protein levels were detected as measured by Azure TotalStain. For Westerns, N=12 for all treatment groups. Equal numbers of male and female mice were used for GFAP Western. No differences between sexes were detected, and data were combined into the figure panel. ANOVA, Tukey’s post hoc testing. For *ex vivo* slice treatment (panel D), slices were obtained from 6 mice (3M/3F) per treatment group, 4 slice/mouse were used. When no sex difference was detected, data were combined by treatment. T-test. See Table 1 for exact statistical values.

Finally, given the astrocyte-restricted expression of AKT2 (Levenga et al., 2021), we sought to determine if acute nicotine exposure impacted AKT2 activation. To do this, we examined AKT2 phosphorylation at Serine 474, a residue critical to AKT2 activation, and total AKT2 levels (Hoeffer & Klann, 2010; Wong et al., 2020). *Ex vivo* hippocampal brain slices were prepared from nicotine-naive animals and incubated in ACSF as we have done previously (Levenga et al., 2017). *S*lices were then treated with nicotine (100 μM) or vehicle for 30 minutes. Following this treatment, significant increases in pAKT2 were observed (**Figure 4D**, vehicle vs. nicotine, p=0.0337). To our surprise, we also detected differences in total AKT2 levels. After nicotine exposure, total AKT2 levels were significantly reduced compared with controls, p=0.0465. Reduction in total levels was confirmed by normalization to total protein loaded per lane **(Figure 4D)**. See Table 1 for specific statistics. Combined, these data support the idea that AKT2 activation, total AKT2 levels, and astrocytes respond to acute nicotine exposure *ex vivo*.

### Astrocytes show decreased morphological complexity following chronic nicotine exposure but show increased complexity when *Akt2* expression is conditionally removed from astrocytes

We next assessed nicotinic responses in astrocytes after a longer exposure period, more aligned with human consumption. To do this, we tested WT and *Akt2* cKO mice using an established model of chronic nicotine exposure (Mathews & Stitzel, 2019). A *Nestin* promoter-driven Cre mouse line was used to generate brain-specific conditional removal of *Akt2* (*Nes-Akt2* cKO) (Levenga et al., 2017). After confirming no innate effect on astrocytes by the *Nes* driver line (**Supplemental Fig 2C**), WT and *Akt2* cKO were given either 0.2% saccharin or nicotine (200 µg/µL in 0.2% saccharin) for two weeks in their drinking water. In contrast to acute exposure, we found that WT astrocytes in the hippocampus had significantly reduced morphological complexity after chronic exposure to nicotine (**Figure 5A, B;** main effect of Sholl radius: p<0.0001, main effect of group: p<0.0001, interaction: p<0.0001; post hoc testing, WT saccharin vs WT nicotine p<0.0001). Without nicotine exposure, cKO astrocytes were smaller than their WT counterparts (**Figure 5A, B;** post hoc testing, WT Saccharin vs. *Akt2* cKO Saccharin: p<0.0001). After chronic nicotine treatment, *Akt2* cKO astrocytes showed significantly increased complexity, opposite that of WT astrocytes. Of note, nicotine treatment made WT astrocytes morphologically indistinguishable from *Akt2* cKO saccharin astrocytes and *Akt2* cKO nicotine astrocytes morphologically indistinguishable from WT saccharin astrocytes (**Figure 5 B;** post hoc testing, *Akt2* cKO Saccharin vs. *Akt2* cKO Nicotine: p<0.0001; other post hoc testing, WT Saccharin vs. *Akt2* cKO Nicotine: p=0.2070; *Akt2* cKO Saccharin vs. WT Nicotine: p=0.4998). See Table 1 for detailed statistics. Astrocyte area was determined next (**Figure 5C**). We found that GFAP-dependent astrocyte area was reduced with chronic nicotine exposure compared with saccharin in WT mice (**Figure 5D**; interaction, p<0.0001; post hoc testing, WT Saccharin vs. WT Nicotine: p=0.0087). Notably, WT saccharin-treated astrocytes were significantly larger than saccharin-treated *Akt2* cKO astrocytes (post hoc testing, WT Saccharin vs. *Akt2* cKO Saccharin: p=0.0026). However, in contrast to the Sholl data (**Figure 5B**), we did not find that nicotine-treated *Akt2* cKO astrocytes were significantly larger than the *Akt2* cKO saccharin group, (**Figure 5D**; post hoc testing, *Akt2* cKO Nicotine vs. *Akt2* cKO Saccharin: p=0.0688). Although there was a trend toward larger *Akt2* cKO astrocytes following chronic nicotine, the effect of chronic nicotine exposure in *Akt2* cKO mice is more pronounced on morphological complexity.

**Figure 5:**
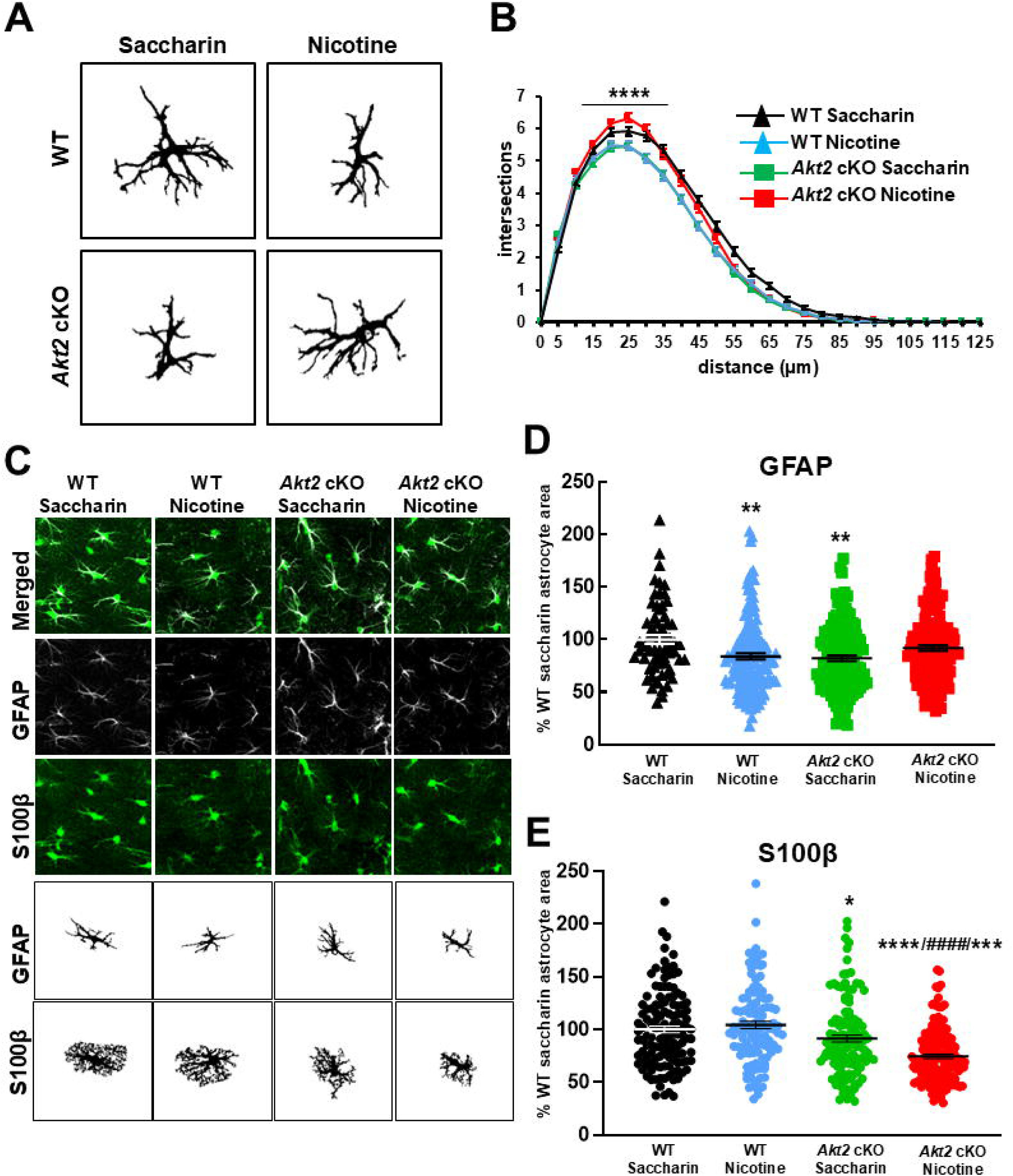
Chronic nicotine treatment results in reduced morphological complexity and GFAP-dependent area in WT astrocytes, and astrocytic AKT2 activity regulates GFAP-dependent and S100β-dependent processes differently. WT and *Nes*-*Akt2* cKO (*Akt2* cKO) mice were exposed to nicotine chronically, and then Sholl analysis was performed on mouse hippocampal astrocytes following treatment. **(A)** Representative binarized images from astrocytes from each experimental group used for Sholl analysis. **(B)** Sholl quantification of WT and *Akt2* cKO responses to chronic nicotine treatment. Following chronic nicotine, WT astrocyte complexity is decreased. Conversely, *Akt2* cKO complexity is increased. WT Saccharin vs. WT Nicotine, WT Saccharin vs. *Akt2* cKO Saccharin, *Akt2* cKO Saccharin vs. *Akt2* cKO Nicotine: **** p<0.0001. **(C)** Representative images from each experimental treatment used for astrocyte area analyses. *Top:* Merged GFAP (white) and S100β (green) staining images of astrocytes in hippocampal sections, with individual channels below. Binarized images for astrocyte area analysis below confocal images. **(D)** GFAP-dependent area analysis of WT and *Akt2* cKO following chronic nicotine treatment. Nicotine-treated WT astrocytes display decreased GFAP-dependent area compared to WT saccharin-treated controls. Likewise, WT saccharin-treated controls are larger than *Akt2* cKO saccharin-treated astrocytes. Saccharin vs. WT Nicotine, WT Saccharin vs. *Akt2* cKO Saccharin: ** p<0.01; **(E)** S100β-dependent area of WT and *Akt2* cKO astrocytes in response to chronic nicotine treatment. Following chronic nicotine, no increase in WT astrocyte area is observed. Conversely, *Akt2* cKO area is decreased after chronic nicotine compared to all control groups. WT Nicotine vs. *Akt2* cKO Saccharin: * p<0.05; *Akt2* cKO Nicotine vs. *Akt2* cKO Saccharin: *** p<0.001; *Akt2* cKO Nicotine vs. WT Nicotine: **** p<0.0001; WT Nicotine vs. *Akt2* cKO Saccharin: #### p<0.0001. Imaging data is representative of 3 stained slices per mouse and 10-20 astrocytes/mouse. Biological N=3 for each group. Groups were alternated by sex, either 2M, 1F, or 2F, 1M. ANOVA, Tukey post hoc. See Table 1 for exact statistical values.

Because GFAP can only approximate a small fraction of total astrocyte volume (Baldwin, Murai, & Khakh, 2024; Ogata & Kosaka, 2002), we further investigated astrocyte area using another marker for astrocyte volume, S100 calcium-binding protein B (S100β). S100β is a glia-specific protein that, in addition to binding calcium, is secreted and plays roles in cell proliferation and growth (Donato, 2003; Donato et al., 2009; Rodriguez et al., 2023; Shashoua, Hesse, & Moore, 1984). S100β is a small, cytoplasmic protein that labels astrocyte volume to a much greater extent than GFAP (**Figure 5C**). Using S100β to estimate astrocyte area, we assessed the effects of chronic nicotine exposure on astrocytes in WT and *Akt2* cKO mice and found differences with genotype and treatment (**Figure 5E**; main effect of treatment: p=0.0351, main effect of genotype: p<0.0001, interaction: p=0.0004). We found no difference in area between WT saccharin and nicotine treated astrocytes whereas *Akt2* cKO astrocytes displayed a significant reduction in area between saccharin and nicotine treatment groups (**Figure 5E**). Astrocytes in the WT nicotine group also had significantly larger S100β-dependent area than in the *Akt2* cKO nicotine or saccharin groups (**Figure 5E**; post hoc testing, WT Nicotine vs. Saccharin *Akt2* cKO: p=0.0158; *Akt2* cKO Saccharin vs. *Akt2* cKO Nicotine: p=0.0002; WT Nicotine vs *Akt2* cKO Nicotine, WT Nicotine vs *Akt2* cKO Saccharin: p<0.0001). In contrast to the GFAP-dependent area analyses, these S100β-based data show no impact of chronic nicotine on astrocyte area in WT mice while chronic nicotine in *Akt2* cKO mice significantly reduced astrocyte area.

Nicotine consumption was not a factor in these findings. Average daily fluid consumption for all nicotine groups across the two-week period was 6.73 ± 0.376 mL/day and was nearly identical between the WT and cKO groups (**Supplemental Figure 5A**). This equates to an average daily consumption of nicotine at 38.46 mg/kg/day, which is consistent with previous reports where nicotine is the sole fluid source (Pietilä & Ahtee, 2000). Because astrocyte numbers could affect these data, astrocyte cell counts in the imaging area were performed. There were no detectable differences in total astrocyte numbers between treatment groups in the imaged CA1 region (**Supplemental Figure 5B**). As with acute nicotine exposure, the morphometric differences we identified were not due to changes in astrocyte number in the CA1 following the drinking protocol or exposure to nicotine.

### α7 nAChRs and α4β2 activity are required for responses to nicotine in primary cultured astrocytes

In primary cultures of mouse astrocytes, we found that nicotine also induces astrocyte activation, consistent with our results in nicotine-exposed mice and in other *in vitro* studies (Aryal et al., 2021). We determined the minimum acute nicotine concentration required to elicit a response in cultured astrocytes was 100 μM nicotine, detected as an increase in GFAP-dependent astrocyte area (**Figure 6A, B;** main effect of treatment: p=0.0004; post hoc testing, vehicle vs. 100µM Nicotine: p=0.0071; 1µM Nicotine vs. 100µM Nicotine: p=0.0014, 10µM Nicotine vs. 100µM Nicotine: p=0.0019). Next, we tested cultured astrocyte responses under conditions of nAChR blockade. Astrocytes express α7 and α4β2 nicotinic acetylcholine receptor (nAChR) subtypes (Aryal et al., 2021; Gahring, Persiyanov, Dunn, et al., 2004; Gahring, Persiyanov, & Rogers, 2004). To determine if either of these nAChRs was involved in facilitating astrocyte responses to nicotine, we used antagonists for each subtype, methyl lycaconitine (MLA) for α7 nAChR and Dihydro-beta-erythroidine (DHβE) for α4β2 (Drasdo, Caulfield, Bertrand, Bertrand, & Wonnacott, 1992; Opanashuk, Pauly, & Hauser, 2001). We found that astrocytes failed to significantly change morphology, as measured by GFAP-dependent area, in response to 100μM nicotine in the presence of 1 µM MLA (**Figure 6A, C;** main effect of treatment: p=0.0145; post hoc testing, vehicle vs. 100µM: p=0.0258; vehicle + MLA vs. 100µM Nicotine: p=0.0280, Vehicle vs. 100µM Nicotine + MLA: p=0.9724). See Table 1 for exact statistics. When 1 µM DhβE was applied to cultured astrocytes, they similarly showed a reduced response to nicotine (**Figure 6A, D;** main effect of treatment: p=0.0006; post hoc testing, vehicle vs. 100µM nicotine: p=0.0165, vehicle + DhβE vs. 100µM Nicotine: p=0.0003; 100µM Nicotine + DhβE vs. Vehicle: p=0.9386). Finally, we examined nicotine response in *Akt2* KO astrocytes. We found that while *Akt2* KO astrocyte area also increased with 100µM nicotine treatment compared with WT controls, nicotine-treated KO astrocytes did not significantly differ from vehicle-treated KO controls, which contrasted with the increased response seen *in vivo.* (**Figure 6A, E;** main effect of treatment p=0.0007, main effect of genotype p= 0.0369; post hoc testing, WT vehicle vs. *Akt2* KO vehicle: p=0.2558, WT vehicle vs. WT 100µM Nicotine: p=0.0356, WT vehicle vs. *Akt2* KO 100uM Nicotine: p=0.0017, *Akt2* KO vehicle vs. *Akt2* KO 100µM Nicotine: p=0.0969, *Akt2* KO 100µM Nicotine vs. WT 100µM Nicotine: p=0.6305). These data demonstrate that α7 and α4β2 nAChRs play a significant role in the astrocytic responses to nicotine and that AKT2 activity is required for normal primary culture responses to nicotine.

**Figure 6:**
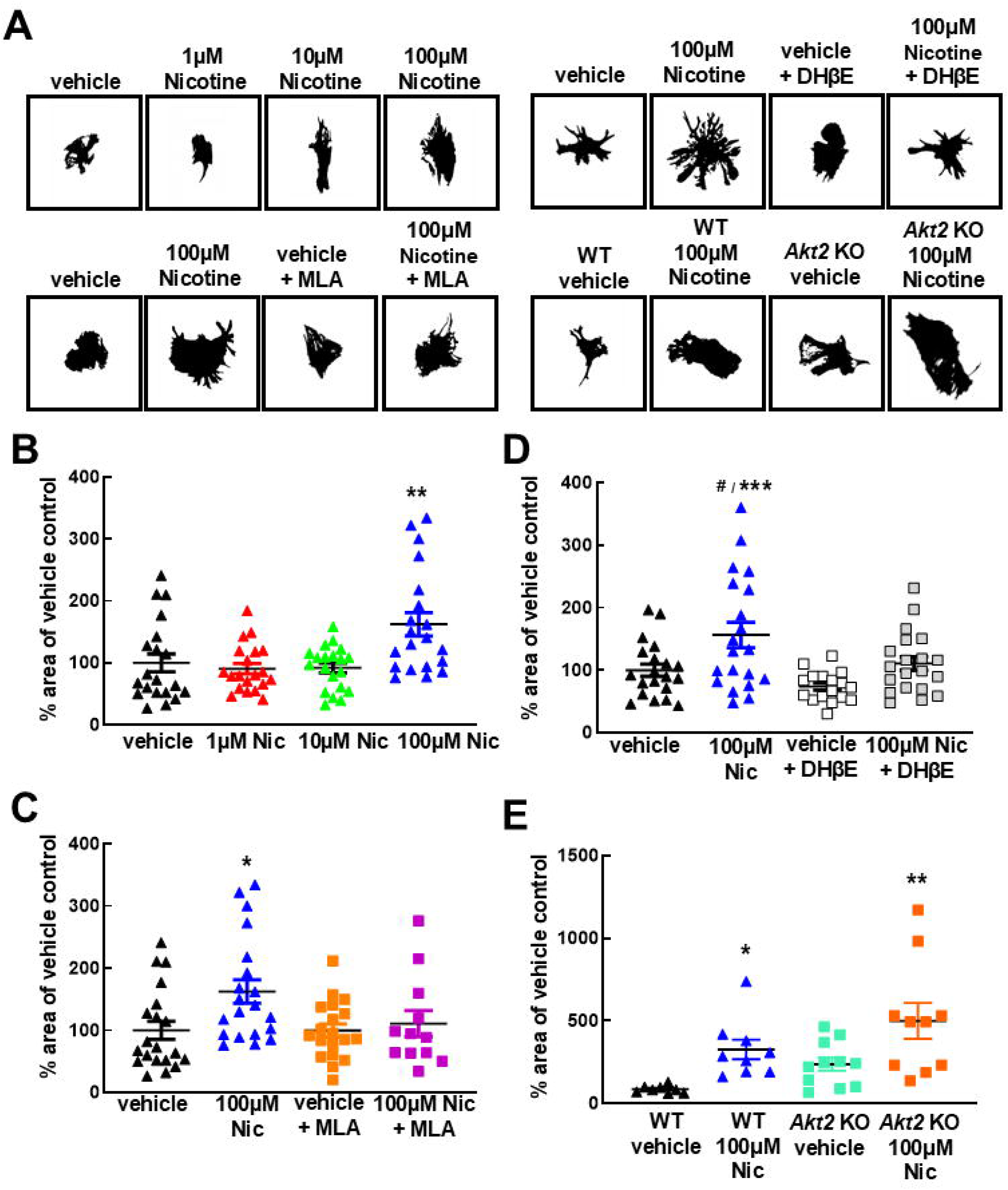
Acute nicotine treatment of cultured astrocytes induces morphological changes consistent with astrocyte activation, mitigated by blocking α7 and α4β2 nicotinic acetylcholine receptors. Primary mouse astrocytes (WT or *Akt2* KO) were cultured and exposed to nicotine under different conditions of nAChR blockade. Area analysis of stained astrocytes was completed on astrocytes processed using standard imaging software. **(A)** Representative binarized images of astrocytes from each experimental group. **(B)** Quantification of astrocyte size following acute nicotine treatment (vehicle, 1µM, 10µM, 100µM). Vehicle vs. 100µM Nicotine, 1µM Nicotine vs. 100µM Nicotine, 10µM Nicotine vs. 100µM Nicotine: **p<0.01. **(C)** Quantification of astrocyte size following acute nicotine treatment (100µM) with or without methyl lycaconitine (MLA) blockade of nAChR (1μM). vehicle vs. 100µM Nicotine, vehicle + MLA vs. 100µM Nicotine: * p<0.05). **(D)** Quantification of astrocyte size following acute nicotine treatment (100µM) with and without Dihydro-beta-erythroidine (DHβE) treatment (1μM). Vehicle vs. 100µM Nicotine: # p<0.05; vehicle + DhβE vs. 100µM Nicotine: ***p<0.001. **(E)** Quantification of astrocyte area following acute nicotine treatment of *Akt2* KO primary astrocytes. Unlike WT astrocytes, *Akt2* KO astrocytes did not show significant increases in area after nicotine exposure. Vehicle-treated *Akt2* KO astrocytes were not significantly larger than vehicle-treated WT astrocytes. WT vehicle vs. WT 100µM Nicotine: * p<0.05; WT vehicle vs. *Akt2* KO 100µM Nicotine ** p<0.01; *Akt2* KO vehicle vs. *Akt2* KO 100µM Nicotine p>0.05; WT vehicle vs. *Akt2* KO vehicle p>0.05. ANOVA, Tukey post hoc. See Table 1 for exact statistics. Data from three independent cultures using 3-5 cells/coverslip for each panel.

### *Akt2* cKO mice showed reduced conditioned place preference (CPP) performance compared with WT littermate controls

After showing that AKT2 is involved in astrocytic nicotine responses *in vivo* and *in vitro* **(Figure 2,4**), we sought to determine if astrocytic AKT2 activity regulated functional outcomes linked to nicotine exposure. Since we observed the acute effects of nicotine treatment on astrocyte activation, we chose to examine the potential role of astrocytic AKT2 in the rewarding effects of nicotine. Because our experiments centered on hippocampal astrocytes, we selected the conditioned place preference (CPP) assay to examine functional outcomes related to AKT2 activity and nicotine. The CPP assay depends on hippocampal function (Ferbinteanu & McDonald, 2001; Hitchcock & Lattal, 2018) and is frequently employed in rodents to test the rewarding effects of stimulus on behavior, which in this case is the preference for a chamber based on its association with nicotine exposure. To test whether astrocyte-specific AKT2 deficiency in *Gfap*-*Akt2* cKO (*Akt2* cKO) mice altered CPP performance in response to a rewarding dose of nicotine, mice were trained to associate a location with nicotine over two conditioning periods (**Figure 7A)** in a three-chambered CPP apparatus (**Figure 7B**). Following one day of preconditioning, mice were conditioned with two daily injections three days in a row before a testing day for each conditioning period. We found that WT mice displayed a significantly higher preference for the chamber in which nicotine was administered compared with *Akt2* cKO mice (**Figure 7C**; p=0.0130). These preference differences were not due to general locomotor responses in either distance traveled (**Figure 7D**; p=0.4100) or in time spent in the unpreferred chamber (exploration) on testing day (**Figure 7E**; p=0.660). See Table 1 for exact statistics. These data implicate AKT2 in astrocytic processes underlying fundamental cognition related to the rewarding effects of nicotine during CPP.

**Figure 7:**
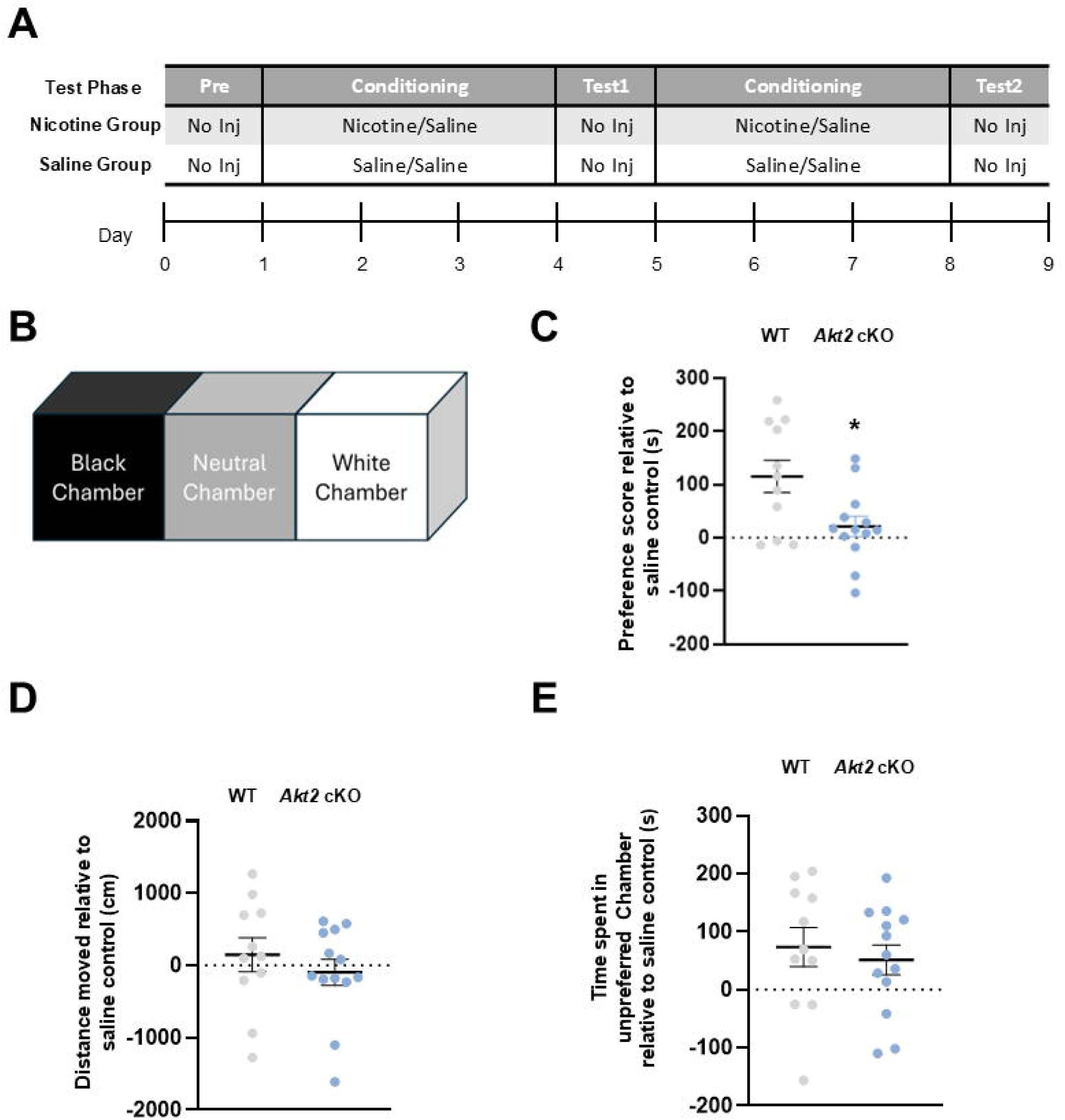
*Akt2* cKO mice show impaired Conditioned Place Preference (CPP). CPP was performed on two cohorts of WT and GFAP-*Akt2* cKO (*Akt2* cKO) mice. **(A)** Preconditioning Day (day 1 of the timeline), injection days (days 2-4 of the timeline), Testing D1 (day 5 of the timeline), injection days (days 6-8 of the timeline), Testing D2 (day 9 of the timeline). **(B)** The three-chamber apparatus used for CPP. The two conditioning chambers and the neutral release chamber are identically sized. **(C)** Data represented as average day 2 preference scores (in seconds) subtracted from saline controls both testing days following CPP training. Significant differences were seen between nicotine-treated *Akt2* cKO and WT mice: * p<.05. **(D)** Distance traveled during testing day 2 relative to saline controls. No differences were detected between genotypes between nicotine and saline groups. **(E)** Time spent in the unpreferred chamber during testing day 2 relative to saline controls. No differences were detected between either genotypes or nicotine and saline groups. 11 WT Nicotine, 13 *Akt2* cKO Nicotine. 12 mice per genotype were used to establish saline preference. Data was collected from three independent behavioral cohorts for each genotype. Sex/Group: Nicotine: WT 6M, 5F; Nicotine *Akt2* cKO 6M, 7F. Saline groups split evenly between male and female mice. T-test See Table 1 for exact statistics.

### Nicotine Alters Hippocampal Transcriptomic Signatures Linked to Smoking Initiation and Neurotransmission

Nicotine induces transcriptomic changes in the hippocampus of genes associated with smoking initiation, neurotransmitter receptor activity, and transmembrane transporter activity. We also examined whether transcriptomic changes occurred to complement the morphological changes we observed in astrocytes in response to nicotine exposure. We performed RNA sequencing on the hippocampal tissue collected 24 hours after nicotine (0.2 mg/kg) versus saline injection (**Figures 3, 4**). We then characterized transcriptional differences with a volcano plot and gene ontology enrichment diagrams (**Figure 8A-C**). Following nicotine treatment, we found 35 differentially expressed genes, with 17 genes upregulated and 18 genes downregulated (**Supplementary Table 2**). A gene ontology over-representation test identified biochemical pathways overrepresented with nicotine exposure. Several pathways related to neurotransmitter receptor activity, such as acetylcholine and serotonin receptor activity, were enriched, likely due to the importance of serotonergic and dopaminergic neurons in nicotine addiction (Zeid, Kutlu, & Gould, 2018). Transmembrane transporter activity was also identified and clustered together using an enrichment map (**Figure 8B**) (G. Yu, L. G. Wang, Y. Han, & Q. Y. He, 2012). Further gene ontology (GO) pathway analysis using Enrichr showed pathways involved in energy metabolism based on GO: Biologic process (E. Y. Chen et al., 2013; Kuleshov et al., 2016; Xie et al., 2021) (**Figure 8C**). Interestingly, structural alterations in mitochondria have been found to occur during the progression of nicotine self-administration (Fan et al., 2024). Other pathways using different search terms, such as smoking initiation, were enriched based on the GWAS Catalog 2023 gene sets (**Figure 8D**). Genes of interest associated with smoking initiation are listed, showing that 24 hours post-nicotine exposure of one dose was sufficient to induce transcriptomic changes in the hippocampus of genes that have already been identified to play a role in smoking initiation according to GWAS studies (**Figure 8E**). Changes to astrocytic morphology may affect astrocytic homeostatic function (PMID: 38943350, 23243501). Some genes involved in astrocytic homeostatic function were found to be differentially expressed in response to nicotine (**Figure 8F**). When compared to LPS, most differed in either magnitude of expression or direction of change compared to LPS, demonstrating that LPS-induced astrogliosis and nicotine-induced astrocyte activation can be differentiated at a homeostatic level (**Figure 8F**). Taken together, these results demonstrate that nicotine can induce changes at the transcriptome level in the mouse hippocampus, and the associated pathways are in overlap with addiction transcriptomic studies in the ventral tegmental area (VTA) and are involved in transmembrane transporter activity, neurotransmitter receptor activity, and smoking initiation (Fan et al., 2024).

**Figure 8:**
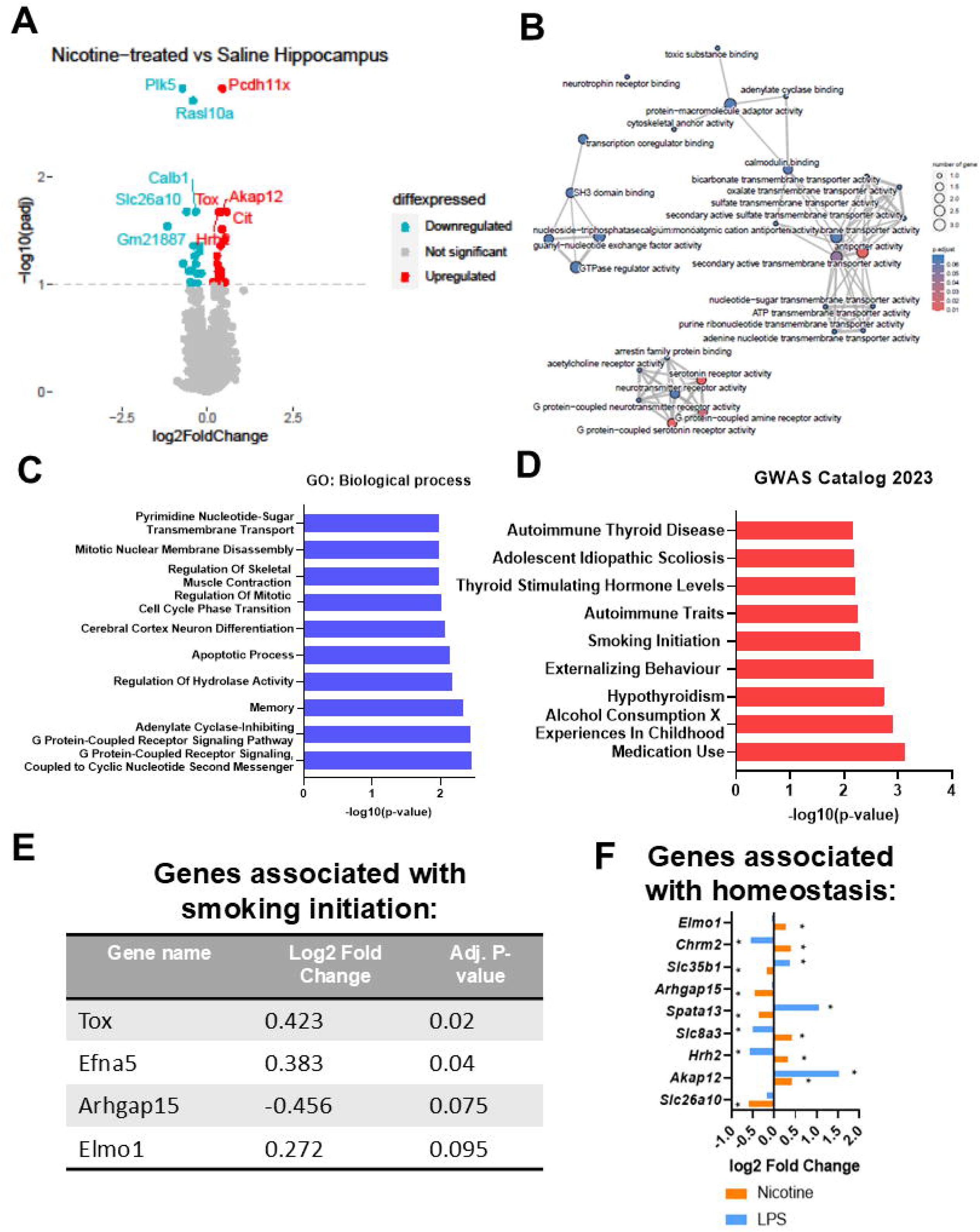
Transcriptomic analyses of hippocampal gene expression 24 hours after in vivo nicotine exposure. (**A**) Volcano plot of significantly differentially expressed genes identified between Nicotine-treated hippocampus compared to control Saline-treated hippocampus. Red dots indicate genes found upregulated and green indicates downregulated genes with an adjusted p-value of 0.1. (**B**) Enrichment map of enriched terms found using gene set enrichment analysis (GSEA) in a network with edges connecting overlapping gene sets. Mutually overlapping gene sets are clustered together. (**C**) Gene Ontology (GO) analysis of significantly differentially expressed genes using Enrichr GO: Biological process or (**D**) GWAS catalog 2023 terms. All analyses are based on RNA-seq data from N=6 WT male mice/group. (**E)** Log2 Fold change and adjusted p values for genes differentially expressed in Smoking Initiation GWAS catalog term. (**F**) Log2 Fold change for genes involved in homeostatic regulation for Nicotine and LPS treatment. * p adj < 0.1. 6 mice per treatment used for RNA preparation and library construction. See Table 1 for specific statistics.

## Discussion

Our study provides new evidence for AKT2’s role in intracellular signaling underlying astrocyte responses to nicotine. We also provide the first evidence for an *in vivo* response to acute nicotine exposure and identify AKT2-dependent modification of astroglial morphological processes linked to nicotine-mediated astrocyte activation. We also identify different roles for AKT2 function depending on the nature of nicotine exposure and cellular context. Consistent with a previous report, we show that nicotine acts through astrocytically expressed α7 nAChRs (Aryal et al., 2021). Adding to this finding, we found *in vitro* that either α7 or α4β2 receptors can mediate astrocyte responses to nicotine. There is some evidence of α7β2 receptors in astrocytes, which could also explain the effects of both antagonists (Wu et al., 2016). In contrast to previous work, we provide evidence of reduced astrocyte morphological responses following *in vivo* chronic nicotine exposure. Our data links astrocytic AKT2 activity to nicotine-CPP behavior. Combined, these data indicate a potential role for astrocytes in nicotine use disorders and a critical role for AKT2 in the underlying neurobiology of nicotine use. More broadly related to astroglial biology, we identify AKT2 as a novel molecular actor in mediating morphological responses to different astrocytic stimuli and suggest that AKT2 is involved in multiple activation-modulation, and cell growth.

These studies provide important new mechanistic insights into underlying intracellular regulation of astrocytic responses to stimuli like nicotine and LPS. Our study is the first to report on the effects of *Akt2* deficiency on the astrocytic response to nicotine or LPS (**Figures 2, 5, 6, 7**), and the activation of AKT2 by nicotine in the brain (**Figure 4**). These data are of primary importance for astroglial biology because of the highly specific expression of AKT2 within CNS astrocytes. Our published research established that AKT isoforms exhibit restricted expression at the protein level in different brain cell types (Levenga et al., 2017; Levenga et al., 2021). AKT2 is only expressed in astrocytes and not expressed in neurons or microglia of mouse and human brains (Levenga et al., 2021). Neuroprotective effects of nAChRs are known to require AKT signaling (Kihara et al., 2001; Kume & Takada-Takatori, 2018; Shaw, Bencherif, & Marrero, 2002), but AKT isoform-dependent signaling has not been studied in nicotine use. AKT2 is expressed elsewhere in the body, but because our study used conditional genetic *Akt2* removal of AKT2, the potential impact of non-CNS AKT2 deficiency on our results is unlikely.

*In vitro* primary astrocyte culture and *in vivo* studies indicate that AKT2 activity is required for normal responses to acute applications of nicotine. While cultured *Akt2* KO astrocytes do not significantly increase in area 24 hours after nicotine treatment (**Figure 6**), they are on average larger than WT astrocytes, which could occlude astrocytic morphological responses. The absence of significant morphological responses in *Akt2* KO cultured astrocytes to nicotine could indicate a developmental factor that mediates AKT2 activity in astrocyte growth. Another strong possibility is that astrocytic AKT2 is regulated by interaction or co-factors derived from other neural cell types in the CNS. This possibility, supported by *in vivo* data in our study, indicates that astrocytes significantly increase morphological activation *in vivo* in response to nicotine exposure in a time-dependent (**Figure 5**) and dose-dependent fashion (**Figure 3**), displayed by altered astrocyte activation marker expression (**Figure 4**). Critically, *Akt2* cKO mice display enhanced responses to LPS (**Figure 2**). However, following chronic exposure *in vivo*, the effects of nicotine and AKT2 deficiency were more complex. Data from Sholl and GFAP-dependent area analyses show that astrocytes reduced their overall morphological complexity, consistent with astrocytic atrophy (**Figure 5B, D**, (A. Verkhratsky, Rodrigues, Pivoriunas, Zorec, & Semyanov, 2019)). However, the S100β-determined area did not significantly reduce astrocyte size in chronically exposed WT mice (**Figure 5E**). The difference between acute and chronic nicotine exposures could be cell-autonomous signaling changes linked to temporal activation dynamics. Another possibility is that other cell types, like neurons, may alter nAChR stoichiometry expression in response to chronic nicotine exposure over time (Xiao, Zhou, Jiang, & Yin, 2020), influencing AKT2-dependent astrocyte physiology. Future studies using single-cell RNASeq or proteomics, temporally examining nicotine-dependent cell-type responses and in vitro co-culture experiments, may shed light on these possibilities. In contrast, *Akt2* cKO astrocytes displayed increased complexity (**Figure 5B**), but reduced S100β-determined area (**Figure 5E**) in response to nicotine. AKT2 activity negatively regulates astrocyte GFAP-related morphological responses to acute LPS treatment (**Figure 2**) and chronic nicotine (**Figure 5B, D**). However, in a chronic nicotine paradigm, AKT2 promotes changes in astrocyte area (**Figure 5E).** These apparently paradoxical findings could indicate the existence of multiple AKT2 astrocytic signaling pathways that independently regulate different cellular pathways related to cytoskeletal rearrangement and cell growth. We explored this idea using a transcriptomic approach to examine nicotine-dependent gene expression in the hippocampus (**Figure 8**). These expression data verify results from (Aryal et al., 2021), confirming that nicotine does not induce astrocytic marker gene expression changes that align with astrogliosis (**Supplemental Figure 3D**). However, they indicate robust transcriptional impacts on G-protein and second messenger signaling, mitotic control, transmembrane transport, and nuclear membrane assembly (**Figure 8**).

Our gene expression data indicate that although LPS and nicotine-induced astrocyte activation appear broadly similar at the morphological level (**Figures 1** and **3, Supplemental Figure 2B**), homeostatic responses to LPS-induced astrogliosis and nicotine are transcriptionally distinct (**Figure 8**). A more detailed examination of astrocyte homeostatic gene expression changes compared to LPS reveals some potential links between smoking and nicotine-dependent astrocyte function. Significantly, our data show that a single dose of nicotine can increase gene expression of *Chrm2*, a muscarinic receptor, which is associated with nicotine dependence (Mobascher et al., 2010). Other homeostatic genes identified by our study such as *Arhgap15* and *Akap12*, have connections to diseases linked with nicotine-dependence (Oldenburger et al., 2014; Sigurdsson et al., 2017). Another homeostatic gene regulated by nicotine in the mouse hippocampus is *Slc26a10.* A recent study showed that epigenetic regulation of *Slc26a10* can be used to infer smoking habits (Ambroa-Conde; 2024). A caveat of these transcriptional data is that they are derived from complex tissue sources. Future work using purified cell populations or single-cell RNASeq from different regions should be explored to increase sensitivity and cellular resolution of nicotine-dependent gene expression regulation.

Separate from transcriptional influences, another strong candidate signaling pathway modulated by nicotine is the AKT-mechanistic target of rapamycin (mTOR) signal cascade, essential to translational regulation, cell growth, and differentiation (Hoeffer & Klann, 2010). mTOR is activated by reactive astrogliosis (Codeluppi et al., 2009; Jeong, Cho, Lee, Kim, & Kim, 2021) and is required for astroglial proliferation (C.-Y. Li et al., 2015). Therefore, one possibility is that AKT2 regulates the translation of astrocytic mRNAs that, in turn, control morphological and functional responses to nicotine. However, because AKT regulates numerous cellular pathways (Cho et al., 2001; Gai et al., 2015; Wong et al., 2020; J. Zhang et al., 2010). Future studies using unbiased genomics and proteomics approaches will be needed to rigorously identify AKT-dependent pathways that manifest astrocyte morphological alterations following nicotine exposure.

Our data suggests that chronic nicotine exposure results in astroglial asthenia. Such a process would reduce the homeostatic supportive and protective functions astroglia provide to neurons and may contribute to an altered pattern of synaptic transmission associated with chronic nicotine exposure (Fujii, Ji, & Sumikawa, 2000; Grieder et al., 2014; Miura, Ishii, Aosaki, & Sumikawa, 2006; Welsby, Rowan, & Anwyl, 2006). It is important to note that our results with chronic nicotine exposure *in vivo* in the hippocampus differ from those reported by Aryal and colleagues, who reported increased astrocyte volume following chronic nicotine (Aryal et al., 2021). Our hippocampal chronic nicotine results derived from Sholl and GFAP- and S100β-dependent area analyses do not align with increased astrocyte area (**Figure 5**). There are a few potential reasons for this difference. First, our astrocyte measures differed. Aryal et al. used *Aldh1l1* driven tdTomato-dependent 3D reconstruction of confocal image stacks for volume analysis from cleared tissues (Aryal et al., 2021) and we examined Z-stack (GFAP or S100β) staining from individual astrocytes. These differ in at least two significant ways. Our tissues underwent disparate processing for astrocyte marker detection and subsequent image analysis. Moreover, we might not have sampled the same astrocyte populations based on small but significant differences in astrocyte *Aldh1l1, GFAP,* and *S100*β gene expression, protein expression, or stability. Second, we delivered our nicotine via drinking water, not through osmotic minipumps. Each approach has advantages and disadvantages, but nicotine delivery in drinking water is well known to provide simple, chronic nicotine exposure and the induction of dependence (Matta et al., 2007). Our primary motivation to use drinking water for oral nicotine instead of minipumps was concern over the potential effects of surgery on astroglial activation in the brain. In other AKT2-related studies by our group, we have found that AKT2 can be activated by inflammatory events such as intraventricular cannula implantation surgery or viral infection (data not shown). We have not assessed the impacts of subcutaneous mini-pump implantation on AKT2, used in (Aryal et al., 2021), so we cannot affirm this to be a critical factor in the differences between our findings. Finally, there are differences in the amount of nicotine delivered for each chronic exposure. Our drinking water dose (200 ug/ml for 14 days) in C57BL/6J mice generates an average blood serum concentration of ∼16 ng/ml (Grabus et al., 2005). The dose used for minipump infusion in the Aryal study was set at 2 mg/kg/h for 12 days, approximating a nicotine plasma concentration of ∼100 ng/mL (Matta et al., 2007). Therefore, it may be that our observed chronic nicotine effects differ from those of Aryal et al. because of lower overall nicotine exposure in our study. Astrocytes may respond differently to a range of nicotine concentrations and exposure times. Future studies would benefit from simultaneously analyzing multiple chronic nicotine concentrations and comparing methods.

Astrocytes express α4β2 and α7 nAChRs (Gahring, Persiyanov, Dunn, et al., 2004; Kume & Takada-Takatori, 2018; Q. Liu et al., 2018; Loram et al., 2012; Patti et al., 2007; Pavlov, Wang, Czura, Friedman, & Tracey, 2003; Sharma & Vijayaraghavan, 2001). Several lines of study show that α4β2 nAChRs mediate many of the effects of acute nicotine on mouse behavior, including learning and memory (Davis & Gould, 2006, 2007; Kutlu, Holliday, & Gould, 2016; Marubio et al., 2003). In rodents, α7 nAChRs are involved in attention and working memory tasks (Young et al., 2004), spatial discrimination (Levin et al., 2009), immune modulation, inflammation (Kume & Takada-Takatori, 2018; Q. Liu et al., 2018; Revathikumar et al., 2016), and neuropathic pain responses (Loram et al., 2012). More recently, the direct involvement of astrocytic α7 nAChRs in associative fear memory was shown (K. Zhang et al., 2021). Our *in vitro* data suggests that α7- and α4β2-containing nAChRs can mediate normal astrocytic nicotine responses (**Figure 6**). One critical limitation of our nAChR blockade experiment is that they were performed *in vitro*. Future work to verify our *in vitro* findings will be essential to validate the nAChRs responsible for mediating nicotine effects on astrocyte populations *in vivo*. Our data does not rule out the possibility of indirect contributions from nAChRs in neuronal populations. They also do not rule out the possibility of more complex interactions between astroglial nAChRs during more long-term nicotine exposure or during development. More detailed analyses using astrocyte-specific removal or manipulation of astrocytically expressed nAChRs will be required to address the contributions of astrocyte nAChRs. This could be achieved using genetic or viral reagents, such as conditional removal of *Akt2* from astrocytes pre- or post-developmentally. Identifying the nAChRs responsible for nicotine responsiveness may be critical in determining the signaling cascades recruited or modified during the distinct phases of nicotine exposure.

Nicotine’s potent effect on cognition, attention, and memory is believed to underlie the formation of strong drug-context associations, which can drive future nicotine use triggered by contextual or environmental cues (Davis, James, Siegel, & Gould, 2005; Kutlu et al., 2016). In this way, the acute effects of nicotine can promote drug-seeking behavior and escalation to chronic use driven by pathological memory formation (Markou, 2008; Markou & Paterson, 2001; Paterson & Markou, 2004). The relationship between contextual cognition and pathological memory formation was our primary motivation for examining hippocampal astrocytes. Therefore, an improved understanding of how nicotine affects cognition, and memory may be critical for preventing nicotine dependence. The CPP paradigm is a classic test that assesses the rewarding properties of a drug and relies on hippocampal function (Ferbinteanu & McDonald, 2001; Hitchcock & Lattal, 2018). Nicotine CPP shows a typical inverted dose-response curve; low doses are without effect, intermediate doses tend to be rewarding, and high doses either are not rewarding or can be aversive. Our experiments utilized an intermediate dose of nicotine to determine if AKT2 was involved in mediating the rewarding effects of nicotine. Our results in WT mice (**Figure 7**) are consistent with previous work on CPP (M. Kutlu, L. Ortega, & T. Gould, 2015; J. Liu et al., 2019). Interestingly, we saw a reduced preference in conditional *Akt2* KO mice compared to controls, consistent with recent findings that AKT isoforms play distinct roles in mouse behaviors such as spatial memory, contextual memory, and extinction (Wong et al., 2020). While the effect of astrocytic AKT2 deficiency did not eliminate CPP, these findings do support a role for AKT2 in signaling the acute rewarding effects of nicotine. Furthermore, nicotine exposure influenced pathways related to neurotransmitter regulation (**Figure 8**), a key component of synaptic plasticity, suggesting a potential role in modulating neuronal signaling, a key part of behavior. Follow-up studies using single-cell approaches with AKT2-deficient mice may be needed to deepen our understanding of the interplay between AKT2 and nicotine in behavior.

Given the importance of understanding molecular mechanisms underlying nicotine reward, future studies should be conducted to examine the potential role of AKT2 in other nicotine-dependent processes. Cognitive impairment is a common symptom during nicotine abstinence (Ashare, Falcone, & Lerman, 2014) and another withdrawal symptom predictive of relapse (Patterson et al., 2010; Rukstalis, Jepson, Patterson, & Lerman, 2005). Working memory is also affected during nicotine withdrawal in humans (Hindocha et al., 2018; Nardone, Shahid, Strasser, Dempsey, & Benowitz, 2020; Sweet et al., 2010) and rodents (Arthur & Levin, 2002; Levin et al., 1990; Levin et al., 2009). Testing whether AKT2 contributes to withdrawal-induced working memory impairments is possible using working memory assays like the radial arm maze task (Pick & Yanai, 1983; Wincott et al., 2014). In summary, our work identifying the role of *Akt2* in nicotine-dependent cognitive effects could be helpful and provide a future therapeutic target for drug abuse and addiction.

## Conclusions

In addition to neurons, astrocytes affect substance use disorders (SUDs) (Alexei Verkhratsky, Steardo, Parpura, & Montana, 2016). We provide *in vivo* data demonstrating that astrocytes respond to nicotine exposure in a time- and dose-dependent manner. Additionally, genome-wide association studies (GWAS) have shown that genetics plays a role in SUDs (Hancock, Markunas, Bierut, & Johnson, 2018), and a TWAS study showed that AKT2 is linked to nicotine use (Saunders et al., 2022). Thus, our study examining AKT2 in astrocyte responses to nicotine provides a critical mechanistic link between genetics and astrocyte function during nicotine dependence. AKT2 has been linked genetically to pathological astroglial activity (Tamim et al., 2014), and now, in this study, it is a loss of function conditional *Akt2* deletion from astrocytes. Previous studies have shown that AKT is involved in the physiological mechanisms of addiction (Zhu et al., 2021) and plays a role in physiological responses to nicotine (Jia et al., 2016; Y. Li et al., 2019; Xu, Ni, Chen, & Dai, 2019). Together, these data address a gap in our understanding of the role of AKT2 in nicotine use and astrocytes. While an essential first step, future studies will be needed to understand further the role of AKT2 in astrogliosis, nicotine exposure, and cognitive processes impacting nicotine use and withdrawal. This knowledge may allow for developing AKT2-related treatments for future therapeutic strategies to alleviate and prevent nicotine use disorders.

## Acknowledgments

These studies were supported by grants from the National Institutes of Health (R01 NS086933-01, R01 AG064465, R01 AG083268, T32 MH016880, R21/R33 DA055781, and T32 AG052371) and funds from the Linda Crnic Institute. Immunofluorescence imaging was performed at the BioFrontiers Institute’s Advanced Light Microscopy Core (RRID: SCR_018302). Laser scanning confocal microscopy was performed on a Nikon A1R microscope supported by NIST-CU Cooperative Agreement award number 70NANB15H226. We thank Dr. Hunter Mathews, Jaeson Chin, Sam Bingham, Andrew Cooper-Sansone, and Luke Link for their technical contributions to this work and Dr. Christopher Link’s useful comments and discussions in preparing the manuscript. We dedicate this work to the loving memory of our friend and colleague, Ms. Lauren LaPlante, whose contributions were critical to this study and the well-being of our research group.

## Notes

### Competing Interest Statement

The authors have declared no competing interest.

### Summary of Updates

Based on the review from GLIA, we have modified the text to make the discussion more concise, added additional control Figures to corroborate measurements of astrocyte area, imaging controls, and genetic controls. Finally, we have added transcriptomic data to improve the mechanistic analysis of study.

## References

Abbondanza, A., Urushadze, A., Alves-Barboza, A. R., & Janickova, H. (2024). Expression and function of nicotinic acetylcholine receptors in specific neuronal populations: Focus on striatal and prefrontal circuits. Pharmacol Res, 204, 107190. doi:10.1016/j.phrs.2024.107190

Allen, S. P., Seehra, R. S., Heath, P. R., Hall, B. P. C., Bates, J., Garwood, C. J., … Simpson, J. E. (2020). Transcriptomic Analysis of Human Astrocytes In Vitro Reveals Hypoxia-Induced Mitochondrial Dysfunction, Modulation of Metabolism, and Dysregulation of the Immune Response. International Journal of Molecular Sciences, 21(21), 8028. doi:10.3390/ijms21218028

Anderson, W., Greenhalgh, A., Takwale, A., David, S., & Vadigepalli, R. (2017). Novel Influences of IL-10 on CNS Inflammation Revealed by Integrated Analyses of Cytokine Networks and Microglial Morphology. Front Cell Neurosci, 11, 233. doi:10.3389/fncel.2017.00233

Araque, A., Martín, E. D., Perea, G., Arellano, J. I., & Buño, W. (2002). Synaptically released acetylcholine evokes Ca2+ elevations in astrocytes in hippocampal slices. The Journal of Neuroscience: The Official Journal of the Society for Neuroscience, 22(7), 2443–2450. doi:10.1523/JNEUROSCI.22-07-02443.2002

Arthur, D., & Levin, E. (2002). Chronic inhibition of α4β2 nicotinic receptors in the ventral hippocampus of rats: impacts on memory and nicotine response. Psychopharmacology, 160(2), 140–145. doi:10.1007/s00213-001-0961-6

Aryal, S. P., Fu, X., Sandin, J. N., Neupane, K. R., Lakes, J. E., Grady, M. E., & Richards, C. I. (2021). Nicotine induces morphological and functional changes in astrocytes via nicotinic receptor activity. Glia, 69(8), 2037–2053. doi:10.1002/glia.24011

Ashare, R. L., Falcone, M., & Lerman, C. (2014). Cognitive function during nicotine withdrawal: Implications for nicotine dependence treatment. Neuropharmacology, 76, 581–591. doi:10.1016/j.neuropharm.2013.04.034

Bakshi, M. V., Azimzadeh, O., Merl-Pham, J., Verreet, T., Hauck, S. M., Benotmane, M. A., … Tapio, S. (2016). In-Utero Low-Dose Irradiation Leads to Persistent Alterations in the Mouse Heart Proteome. PLoS One, 11(6), e0156952. doi:10.1371/journal.pone.0156952

Baldwin, K. T., Murai, K. K., & Khakh, B. S. (2024). Astrocyte morphology. Trends Cell Biol, 34(7), 547–565. doi:10.1016/j.tcb.2023.09.006

Barres, B. A. (2008). The mystery and magic of glia: a perspective on their roles in health and disease. Neuron, 60(3), 430–440. doi:10.1016/j.neuron.2008.10.013

Batiuk, M. Y., Martirosyan, A., Wahis, J., de Vin, F., Marneffe, C., Kusserow, C., … Holt, M. G. (2020). Identification of region-specific astrocyte subtypes at single cell resolution. Nat Commun, 11(1), 1220. doi:10.1038/s41467-019-14198-8

Chellian, R., Behnood-Rod, A., Bruijnzeel, D. M., Wilson, R., Pandy, V., & Bruijnzeel, A. W. (2021). Rodent models for nicotine withdrawal. Journal of Psychopharmacology (Oxford, England), 35(10), 1169–1187. doi:10.1177/02698811211005629

Chen, E. Y., Tan, C. M., Kou, Y., Duan, Q., Wang, Z., Meirelles, G. V., … Ma’ayan, A. (2013). Enrichr: interactive and collaborative HTML5 gene list enrichment analysis tool. BMC Bioinformatics, 14, 128. doi:10.1186/1471-2105-14-128

Chen, S., Zhou, Y., & Chen, Y. (2018). Jia Gu, fastp: an ultra-fast all-in-one FASTQ preprocessor. Bioinformatics, 34(ue 17). doi:10.1093/bioinformatics/bty560

Cho, H., Mu, J., Kim, J. K., Thorvaldsen, J. L., Chu, Q., Crenshaw, E. B., … Birnbaum, M. J. (2001). Insulin Resistance and a Diabetes Mellitus-Like Syndrome in Mice Lacking the Protein Kinase Akt2 (PKBβ). Science, 292(5522), 1728–1731. doi:10.1126/science.292.5522.1728

Codeluppi, S., Svensson, C. I., Hefferan, M. P., Valencia, F., Silldorff, M. D., Oshiro, M., … Pasquale, E. B. (2009). The Rheb–mTOR Pathway Is Upregulated in Reactive Astrocytes of the Injured Spinal Cord. The Journal of Neuroscience, 29(4), 1093–1104. doi:10.1523/JNEUROSCI.4103-08.2009

Corkrum, M., Rothwell, P. E., Thomas, M. J., Kofuji, P., & Araque, A. (2019). Opioid-Mediated Astrocyte-Neuron Signaling in the Nucleus Accumbens. Cells, 8(6). doi:10.3390/cells8060586

Davis, J. A., & Gould, T. J. (2006). The effects of DHBE and MLA on nicotine-induced enhancement of contextual fear conditioning in C57BL/6 mice. Psychopharmacology, 184(3-4), 345–352. doi:10.1007/s00213-005-0047-y

Davis, J. A., & Gould, T. J. (2007). β2 subunit-containing nicotinic receptors mediate the enhancing effect of nicotine on trace cued fear conditioning in C57BL/6 mice. Psychopharmacology, 190(3), 343–352. doi:10.1007/s00213-006-0624-8

Davis, J. A., James, J. R., Siegel, S. J., & Gould, T. J. (2005). Withdrawal from Chronic Nicotine Administration Impairs Contextual Fear Conditioning in C57BL/6 Mice. The Journal of Neuroscience, 25(38), 8708–8713. doi:10.1523/JNEUROSCI.2853-05.2005

Delbro, D., Westerlund, A., Bjorklund, U., & Hansson, E. (2009). In inflammatory reactive astrocytes co-cultured with brain endothelial cells nicotine-evoked Ca(2+) transients are attenuated due to interleukin-1beta release and rearrangement of actin filaments. Neuroscience, 159(2), 770–779. doi:10.1016/j.neuroscience.2009.01.005

Ding, Z. B., Song, L. J., Wang, Q., Kumar, G., Yan, Y. Q., & Ma, C. G. (2021). Astrocytes: a double-edged sword in neurodegenerative diseases. Neural Regen Res, 16(9), 1702–1710. doi:10.4103/1673-5374.306064

Dobin, A., Davis, C. A., Schlesinger, F., Drenkow, J., Zaleski, C., Jha, S., … Gingeras, T. R. (2013). STAR: ultrafast universal RNA-seq aligner. Bioinformatics, 29(1), 15–21. doi:10.1093/bioinformatics/bts635

Donato, R. (2003). Intracellular and extracellular roles of S100 proteins. Microsc Res Tech, 60(6), 540–551. doi:10.1002/jemt.10296

Donato, R., Sorci, G., Riuzzi, F., Arcuri, C., Bianchi, R., Brozzi, F., … Giambanco, I. (2009). S100B’s double life: intracellular regulator and extracellular signal. Biochim Biophys Acta, 1793(6), 1008–1022. doi:10.1016/j.bbamcr.2008.11.009

Drasdo, A., Caulfield, M., Bertrand, D., Bertrand, S., & Wonnacott, S. (1992). Methyl lycaconitine: A novel nicotinic antagonist. Molecular and Cellular Neurosciences, 3(3), 237–243. doi:10.1016/1044-7431(92)90043-2

Ebert, T., Heinz, D. E., Almeida-Correa, S., Cruz, R., Dethloff, F., Stark, T., … Wotjak, C. T. (2021). Myo-Inositol Levels in the Dorsal Hippocampus Serve as Glial Prognostic Marker of Mild Cognitive Impairment in Mice. Front Aging Neurosci, 13, 731603. doi:10.3389/fnagi.2021.731603

Escartin, C., Galea, E., Lakatos, A., O’Callaghan, J. P., Petzold, G. C., Serrano-Pozo, A., … Verkhratsky, A. (2021). Reactive astrocyte nomenclature, definitions, and future directions. Nat Neurosci, 24(3), 312–325. doi:10.1038/s41593-020-00783-4

Fan, L., Liu, B., Yao, R., Gao, X., Wang, H., Jiang, S., … Hu, Q. (2024). Nicotine-induced transcriptional changes and mitochondrial dysfunction in the ventral tegmental area revealed by single-nucleus transcriptomics. J Genet Genomics, 51(11), 1237–1251. doi:10.1016/j.jgg.2024.08.009

Ferbinteanu, J., & McDonald, R. J. (2001). Dorsal/ventral hippocampus, fornix, and conditioned place preference. Hippocampus, 11(2), 187–200. doi:10.1002/hipo.1036

Fujii, S., Ji, Z., & Sumikawa, K. (2000). Inactivation of α7 ACh receptors and activation of non-α7 ACh receptors both contribute to long term potentiation induction in the hippocampal CA1 region. Neuroscience Letters, 286(2), 134–138. doi:10.1016/S0304-3940(00)01076-4

Gahring, L. C., Persiyanov, K., Dunn, D., Weiss, R., Meyer, E. L., & Rogers, S. W. (2004). Mouse strain-specific nicotinic acetylcholine receptor expression by inhibitory interneurons and astrocytes in the dorsal hippocampus. The Journal of Comparative Neurology, 468(3), 334–346. doi:10.1002/cne.10943

Gahring, L. C., Persiyanov, K., & Rogers, S. W. (2004). Neuronal and astrocyte expression of nicotinic receptor subunit beta4 in the adult mouse brain. The Journal of Comparative Neurology, 468(3), 322–333. doi:10.1002/cne.10942

Gai, D., Haan, E., Scholar, M., Nicholl, J., & Yu, S. (2015). Phenotypes of *AKT3* deletion: A case report and literature review. American Journal of Medical Genetics Part A, 167(1), 174–179. doi:10.1002/ajmg.a.36710

Gao, R., Schneider, A. M., Mulloy, S. M., & Lee, A. M. (2024). Expression pattern of nicotinic acetylcholine receptor subunit transcripts in neurons and astrocytes in the ventral tegmental area and locus coeruleus. The European Journal of Neuroscience, 59(9), 2225–2239. doi:10.1111/ejn.16109

Grabus, S. D., Martin, B. R., Batman, A. M., Tyndale, R. F., Sellers, E., & Damaj, M. I. (2005). Nicotine physical dependence and tolerance in the mouse following chronic oral administration. Psychopharmacology, 178(2-3), 183–192. doi:10.1007/s00213-004-2007-3

Grieder, T. E., Herman, M. A., Contet, C., Tan, L. A., Vargas-Perez, H., Cohen, A., … George, O. (2014). VTA CRF neurons mediate the aversive effects of nicotine withdrawal and promote intake escalation. Nature Neuroscience, 17(12), 1751–1758. doi:10.1038/nn.3872

Gusel’nikova, V. V., & Korzhevskiy, D. E. (2015). NeuN As a Neuronal Nuclear Antigen and Neuron Differentiation Marker. Acta Naturae, 7(2), 42–47.

Halassa, M. M., Fellin, T., & Haydon, P. G. (2007). The tripartite synapse: roles for gliotransmission in health and disease. Trends in Molecular Medicine, 13(2), 54–63. doi:10.1016/j.molmed.2006.12.005

Hameed, M. Q., Hui, B., Lin, R., MacMullin, P. C., Pascual-Leone, A., Vermudez, S. A. D., & Rotenberg, A. (2023). Depressed glutamate transporter 1 expression in a mouse model of Dravet syndrome. Ann Clin Transl Neurol, 10(9), 1695–1699. doi:10.1002/acn3.51851

Hancock, D. B., Markunas, C. A., Bierut, L. J., & Johnson, E. O. (2018). Human Genetics of Addiction: New Insights and Future Directions. Current Psychiatry Reports, 20(2), 8. doi:10.1007/s11920-018-0873-3

Hasel, P., Rose, I. V. L., Sadick, J. S., Kim, R. D., & Liddelow, S. A. (2021). Neuroinflammatory astrocyte subtypes in the mouse brain. Nat Neurosci, 24(10), 1475–1487. doi:10.1038/s41593-021-00905-6

Hindocha, C., Freeman, T. P., Grabski, M., Crudgington, H., Davies, A. C., Stroud, J. B., … Curran, H. V. (2018). The effects of cannabidiol on impulsivity and memory during abstinence in cigarette dependent smokers. Scientific Reports, 8(1), 7568. doi:10.1038/s41598-018-25846-2

Hitchcock, L. N., & Lattal, K. M. (2018). Involvement of the dorsal hippocampus in expression and extinction of cocaine-induced conditioned place preference. Hippocampus, 28(3), 226–238. doi:10.1002/hipo.22826

Hoeffer, C. A., & Klann, E. (2010). mTOR signaling: At the crossroads of plasticity, memory and disease. Trends in Neurosciences, 33(2), 67–75. doi:10.1016/j.tins.2009.11.003

Huang, X., Li, X., Shen, H., Zhao, Y., Zhou, Z., Wang, Y., … Qiu, Y. (2023). Transcriptional repression of beige fat innervation via a YAP/TAZ-S100B axis. Nat Commun, 14(1), 7102. doi:10.1038/s41467-023-43021-8

Iacobas, D. A., Iacobas, S., Stout, R. F., & Spray, D. C. (2020). Cellular Environment Remodels the Genomic Fabrics of Functional Pathways in Astrocytes. Genes, 11(5), 520. doi:10.3390/genes11050520

Jagadeeshaprasad, M. G., Govindappa, P. K., Nelson, A. M., Noble, M. D., & Elfar, J. C. (2022). 4-Aminopyridine Induces Nerve Growth Factor to Improve Skin Wound Healing and Tissue Regeneration. Biomedicines, 10(7). doi:10.3390/biomedicines10071649

Jamal, A., King, B. A., Neff, L. J., Whitmill, J., Babb, S. D., & Graffunder, C. M. (2016). Current Cigarette Smoking Among Adults — United States, 2005–2015. MMWR. Morbidity and Mortality Weekly Report, 65(44), 1205–1211. doi:10.15585/mmwr.mm6544a2

Jeong, K. H., Cho, K. O., Lee, M. Y., Kim, S. Y., & Kim, W. J. (2021). Vascular endothelial growth factor receptor-3 regulates astroglial glutamate transporter-1 expression via mTOR activation in reactive astrocytes following pilocarpine-induced status epilepticus. Glia, 69(2), 296–309. doi:10.1002/glia.23897

Ji, R.-R., Donnelly, C. R., & Nedergaard, M. (2019). Astrocytes in chronic pain and itch. Nature Reviews Neuroscience, 20(11), 667–685. doi:10.1038/s41583-019-0218-1

Jia, Y., Sun, H., Wu, H., Zhang, H., Zhang, X., Xiao, D., … Wang, Y. (2016). Nicotine Inhibits Cisplatin-Induced Apoptosis via Regulating α5-nAChR/AKT Signaling in Human Gastric Cancer Cells. PLoS One, 11(2), e0149120. doi:10.1371/journal.pone.0149120

Jurga, A. M., Paleczna, M., Kadluczka, J., & Kuter, K. Z. (2021). Beyond the GFAP-Astrocyte Protein Markers in the Brain. Biomolecules, 11(9). doi:10.3390/biom11091361

Kihara, T., Shimohama, S., Sawada, H., Honda, K., Nakamizo, T., Shibasaki, H., … Akaike, A. (2001). α7 Nicotinic Receptor Transduces Signals to Phosphatidylinositol 3-Kinase to Block A β-Amyloid-induced Neurotoxicity. Journal of Biological Chemistry, 276(17), 13541–13546. doi:10.1074/jbc.M008035200

Kim, H., Huh, Y.-J., Kim, J. H., Jo, M., Shin, J.-H., Park, S. C., … Lee, Y. (2022). Identification and evaluation of midbrain specific longevity-related genes in exceptionally long-lived but healthy mice. Frontiers in Aging Neuroscience, 14, 1030807. doi:10.3389/fnagi.2022.1030807

Knackstedt, L. A., LaRowe, S., Mardikian, P., Malcolm, R., Upadhyaya, H., Hedden, S., … Kalivas, P. W. (2009). The role of cystine-glutamate exchange in nicotine dependence in rats and humans. Biol Psychiatry, 65(10), 841–845. doi:10.1016/j.biopsych.2008.10.040

Kuleshov, M. V., Jones, M. R., Rouillard, A. D., Fernandez, N. F., Duan, Q., Wang, Z., … Ma’ayan, A. (2016). Enrichr: a comprehensive gene set enrichment analysis web server 2016 update. Nucleic Acids Res, 44(W1), W90–97. doi:10.1093/nar/gkw377

Kume, T., & Takada-Takatori, Y. (2018). Nicotinic Acetylcholine Receptor Signaling: Roles in Neuroprotection. In A. Akaike, S. Shimohama, & Y. Misu (Eds.), Nicotinic Acetylcholine Receptor Signaling in Neuroprotection (pp. 59–71). Singapore: Springer Singapore.

Kutlu, M., Ortega, L., & Gould, T. (2015). Strain-dependent performance in nicotine-induced conditioned place preference. Behav Neurosci, 129(1), 37–41. doi:10.1037/bne0000029

Kutlu, M., Ortega, L., & Gould, T. (2015). Strain-dependent performance in nicotine-induced conditioned place preference. Behavioral Neuroscience, 129(1), 37–41. doi:10.1037/bne0000029

Kutlu, M. G., Holliday, E., & Gould, T. (2016). High-affinity α4β2 nicotinic receptors mediate the impairing effects of acute nicotine on contextual fear extinction. Neurobiology of Learning and Memory, 128, 17–22. doi:10.1016/j.nlm.2015.11.021

Levenga, J., Krishnamurthy, P., Rajamohamedsait, H., Wong, H., Franke, T. F., Cain, P., … Hoeffer, C. A. (2013). Tau pathology induces loss of GABAergic interneurons leading to altered synaptic plasticity and behavioral impairments. Acta Neuropathol Commun, 1, 34. doi:10.1186/2051-5960-1-34

Levenga, J., Wong, H., Milstead, R., Keller, B., LaPlante, L., & Hoeffer, C. (2017). AKT isoforms have distinct hippocampal expression and roles in synaptic plasticity. Elife, 6, e30640. doi:10.7554/eLife.30640

Levenga, J., Wong, H., Milstead, R., LaPlante, L., & Hoeffer, C. (2021). Immunohistological Examination of AKT Isoforms in the Brain: Cell-Type Specificity That May Underlie AKT’s Role in Complex Brain Disorders and Neurological Disease. Cerebral Cortex Communications, 2(2), tgab036. doi:10.1093/texcom/tgab036

Levin, E. D., Lee, C., Rose, J. E., Reyes, A., Ellison, G., Jarvik, M., & Gritz, E. (1990). Chronic nicotine and withdrawal effects on radial-arm maze performance in rats. Behavioral and Neural Biology, 53(2), 269–276. doi:10.1016/0163-1047(90)90509-5

Levin, E. D., Petro, A., Rezvani, A. H., Pollard, N., Christopher, N. C., Strauss, M., … Rose, J. E. (2009). Nicotinic α7- or β2-containing receptor knockout: Effects on radial-arm maze learning and long-term nicotine consumption in mice. Behavioural Brain Research, 196(2), 207–213. doi:10.1016/j.bbr.2008.08.048

Li, C.-Y., Li, X., Liu, S.-F., Qu, W.-S., Wang, W., & Tian, D.-S. (2015). Inhibition of mTOR pathway restrains astrocyte proliferation, migration and production of inflammatory mediators after oxygen–glucose deprivation and reoxygenation. Neurochemistry International, 83-84, 9–18. doi:10.1016/j.neuint.2015.03.001

Li, Y., Song, A. M., Fu, Y., Walayat, A., Yang, M., Jian, J., … Xiao, D. (2019). Perinatal nicotine exposure alters Akt/GSK-3β/mTOR/autophagy signaling, leading to development of hypoxic-ischemic-sensitive phenotype in rat neonatal brain. American Journal of Physiology-Regulatory, Integrative and Comparative Physiology, 317(6), R803–R813. doi:10.1152/ajpregu.00218.2019

Liao, Y., Smyth, G. K., & Shi, W. (2014). featureCounts: an efficient general purpose program for assigning sequence reads to genomic features. Bioinformatics, 30(7), 923–930. doi:10.1093/bioinformatics/btt656

Liddelow, S. A., Guttenplan, K. A., Clarke, L. E., Bennett, F. C., Bohlen, C. J., Schirmer, L., … Barres, B. A. (2017). Neurotoxic reactive astrocytes are induced by activated microglia. Nature, 541(7638), 481–487. doi:10.1038/nature21029

Liu, J., Tao, X., Liu, F., Hu, Y., Xue, S., Wang, Q., … Zhang, R. (2019). Behavior and Hippocampal Epac Signaling to Nicotine CPP in Mice. Translational Neuroscience, 10, 254–259. doi:10.1515/tnsci-2019-0041

Liu, Q., Liu, C., Jiang, L., Li, M., Long, T., He, W., … Zhou, J. (2018). α7 Nicotinic acetylcholine receptor-mediated anti-inflammatory effect in a chronic migraine rat model via the attenuation of glial cell activation. Journal of Pain Research, Volume 11, 1129–1140. doi:10.2147/JPR.S159146

Liu, Y., Hu, J., Wu, J., Zhu, C., Hui, Y., Han, Y., … Fan, W. (2012). alpha7 nicotinic acetylcholine receptor-mediated neuroprotection against dopaminergic neuron loss in an MPTP mouse model via inhibition of astrocyte activation. J Neuroinflammation, 9, 98. doi:10.1186/1742-2094-9-98

Loram, L. C., Taylor, F. R., Strand, K. A., Maier, S. F., Speake, J. D., Jordan, K. G., … Watkins, L. R. (2012). Systemic Administration of an Alpha-7 Nicotinic Acetylcholine Agonist Reverses Neuropathic Pain in Male Sprague Dawley Rats. The Journal of Pain, 13(12), 1162–1171. doi:10.1016/j.jpain.2012.08.009

Love, M. I., Huber, W., & Anders, S. (2014). Moderated estimation of fold change and dispersion for RNA-seq data with DESeq2. Genome Biol, 15(12), 550. doi:10.1186/s13059-014-0550-8

Ma, W., Si, T., Wang, Z., Wen, P., Zhu, Z., Liu, Q., … Li, Q. (2023). Astrocytic alpha4-containing nAChR signaling in the hippocampus governs the formation of temporal association memory. Cell Rep, 42(7), 112674. doi:10.1016/j.celrep.2023.112674

Mahmoud, S., Gharagozloo, M., Simard, C., & Gris, D. (2019). Astrocytes Maintain Glutamate Homeostasis in the CNS by Controlling the Balance between Glutamate Uptake and Release. Cells, 8(2), 184. doi:10.3390/cells8020184

Markou, A. (2008). Neurobiology of nicotine dependence. Philosophical Transactions of the Royal Society B: Biological Sciences, 363(1507), 3159–3168. doi:10.1098/rstb.2008.0095

Markou, A., & Paterson, N. E. (2001). The nicotinic antagonist methyllycaconitine has differential effects on nicotine self-administration and nicotine withdrawal in the rat. Nicotine & Tobacco Research, 3(4), 361–373. doi:10.1080/14622200110073380

Marubio, L. M., Gardier, A. M., Durier, S., David, D., Klink, R., Arroyo-Jimenez, M. M., … Changeux, J. P. (2003). Effects of nicotine in the dopaminergic system of mice lacking the alpha4 subunit of neuronal nicotinic acetylcholine receptors. European Journal of Neuroscience, 17(7), 1329–1337. doi:10.1046/j.1460-9568.2003.02564.x

Mathews, H. L., & Stitzel, J. A. (2019). The effects of oral nicotine administration and abstinence on sleep in male C57BL/6J mice. Psychopharmacology, 236(4), 1335–1347. doi:10.1007/s00213-018-5139-6

Matta, S. G., Balfour, D. J., Benowitz, N. L., Boyd, R. T., Buccafusco, J. J., Caggiula, A. R., … Zirger, J. M. (2007). Guidelines on nicotine dose selection for in vivo research. Psychopharmacology, 190(3), 269–319. doi:10.1007/s00213-006-0441-0

Matusova, Z., Hol, E. M., Pekny, M., Kubista, M., & Valihrach, L. (2023). Reactive astrogliosis in the era of single-cell transcriptomics. Front Cell Neurosci, 17, 1173200. doi:10.3389/fncel.2023.1173200

McClure, E. A., Gipson, C. D., Malcolm, R. J., Kalivas, P. W., & Gray, K. M. (2014). Potential role of N-acetylcysteine in the management of substance use disorders. CNS Drugs, 28(2), 95–106. doi:10.1007/s40263-014-0142-x

Miller, K. A., Degan, S., Wang, Y., Cohen, J., Ku, S. Y., Goodrich, D. W., & Gelman, I. H. (2023). PTEN-regulated PI3K-p110 and AKT isoform plasticity controls metastatic prostate cancer progression. Oncogene. doi:10.1038/s41388-023-02875-4

Miura, M., Ishii, K., Aosaki, T., & Sumikawa, K. (2006). Chronic nicotine treatment increases GABAergic input to striatal neurons. NeuroReport, 17(5), 537–540. doi:10.1097/01.wnr.0000204984.21748.e3

Mizrak, D., Levitin, H. M., Delgado, A. C., Crotet, V., Yuan, J., Chaker, Z., … Doetsch, F. (2019). Single-Cell Analysis of Regional Differences in Adult V-SVZ Neural Stem Cell Lineages. Cell Rep, 26(2), 394–406 e395. doi:10.1016/j.celrep.2018.12.044

Mobascher, A., Rujescu, D., Mittelstrass, K., Giegling, I., Lamina, C., Nitz, B., … Winterer, G. (2010). Association of a variant in the muscarinic acetylcholine receptor 2 gene (CHRM2) with nicotine addiction. Am J Med Genet B Neuropsychiatr Genet, 153B(2), 684–690. doi:10.1002/ajmg.b.31011

Nardone, N., Shahid, M., Strasser, A. A., Dempsey, D. A., & Benowitz, N. L. (2020). The influence of nicotine metabolic rate on working memory over 6 hours of abstinence from nicotine. Pharmacology Biochemistry and Behavior, 188, 172836. doi:10.1016/j.pbb.2019.172836

Norden, D. M., Trojanowski, P. J., Villanueva, E., Navarro, E., & Godbout, J. P. (2016). Sequential activation of microglia and astrocyte cytokine expression precedes increased Iba-1 or GFAP immunoreactivity following systemic immune challenge. Glia, 64(2), 300–316. doi:10.1002/glia.22930

Ogata, K., & Kosaka, T. (2002). Structural and quantitative analysis of astrocytes in the mouse hippocampus. Neuroscience, 113(1), 221–233. doi:10.1016/s0306-4522(02)00041-6

Oikawa, H., Nakamichi, N., Kambe, Y., Ogura, M., & Yoneda, Y. (2005). An increase in intracellular free calcium ions by nicotinic acetylcholine receptors in a single cultured rat cortical astrocyte. J Neurosci Res, 79(4), 535–544. doi:10.1002/jnr.20398

Oldenburger, A., Poppinga, W. J., Kos, F., de Bruin, H. G., Rijks, W. F., Heijink, I. H., … Schmidt, M. (2014). A-kinase anchoring proteins contribute to loss of E-cadherin and bronchial epithelial barrier by cigarette smoke. Am J Physiol Cell Physiol, 306(6), C585–597. doi:10.1152/ajpcell.00183.2013

Opanashuk, L. A., Pauly, J. R., & Hauser, K. F. (2001). Effect of nicotine on cerebellar granule neuron development. The European Journal of Neuroscience, 13(1), 48–56.

Patel, H., McIntire, J., Ryan, S., Dunah, A., & Loring, R. (2017). Anti-inflammatory effects of astroglial alpha7 nicotinic acetylcholine receptors are mediated by inhibition of the NF-kappaB pathway and activation of the Nrf2 pathway. J Neuroinflammation, 14(1), 192. doi:10.1186/s12974-017-0967-6

Paterson, N. E., & Markou, A. (2004). Prolonged nicotine dependence associated with extended access to nicotine self-administration in rats. Psychopharmacology, 173(1-2), 64–72. doi:10.1007/s00213-003-1692-7

Patterson, F., Jepson, C., Loughead, J., Perkins, K., Strasser, A. A., Siegel, S., … Lerman, C. (2010). Working memory deficits predict short-term smoking resumption following brief abstinence. Drug and Alcohol Dependence, 106(1), 61–64. doi:10.1016/j.drugalcdep.2009.07.020

Patti, L., Raiteri, L., Grilli, M., Zappettini, S., Bonanno, G., & Marchi, M. (2007). Evidence that α7 nicotinic receptor modulates glutamate release from mouse neocortical gliosomes. Neurochemistry International, 51(1), 1–7. doi:10.1016/j.neuint.2007.03.003

Pavlov, V. A., Wang, H., Czura, C. J., Friedman, S. G., & Tracey, K. J. (2003). The cholinergic anti-inflammatory pathway: a missing link in neuroimmunomodulation. Molecular Medicine (Cambridge, Mass.), 9(5-8), 125–134.

Pekny, M., & Nilsson, M. (2005). Astrocyte activation and reactive gliosis. Glia, 50(4), 427–434. doi:10.1002/glia.20207

Pekny, M., & Pekna, M. (2014). Astrocyte reactivity and reactive astrogliosis: costs and benefits. Physiol Rev, 94(4), 1077–1098. doi:10.1152/physrev.00041.2013

Perea, G., Navarrete, M., & Araque, A. (2009). Tripartite synapses: astrocytes process and control synaptic information. Trends in Neurosciences, 32(8), 421–431. doi:10.1016/j.tins.2009.05.001

Periyasamy, P., Guo, M.-L., & Buch, S. (2016). Cocaine induces astrocytosis through ER stress-mediated activation of autophagy. Autophagy, 12(8), 1310–1329. doi:10.1080/15548627.2016.1183844

Pick, C. G., & Yanai, J. (1983). Eight Arm Maze for Mice. International Journal of Neuroscience, 21(1-2), 63–66. doi:10.3109/00207458308986121

Pietilä, K., & Ahtee, L. (2000). Chronic nicotine administration in the drinking water affects the striatal dopamine in mice. Pharmacology, Biochemistry, and Behavior, 66(1), 95–103. doi:10.1016/s0091-3057(00)00235-5

Preston, A. N., Farr, J. D., O’Neill, B. K., Thompson, K. K., Tsirka, S. E., & Laughlin, S. T. (2018). Visualizing the Brain’s Astrocytes with Diverse Chemical Scaffolds. ACS Chem Biol, 13(6), 1493–1498. doi:10.1021/acschembio.8b00391

Qian, Z., Qin, J., Lai, Y., Zhang, C., & Zhang, X. (2023). Large-Scale Integration of Single-Cell RNA-Seq Data Reveals Astrocyte Diversity and Transcriptomic Modules across Six Central Nervous System Disorders. Biomolecules, 13(4). doi:10.3390/biom13040692

Revathikumar, P., Bergqvist, F., Gopalakrishnan, S., Korotkova, M., Jakobsson, P.-J., Lampa, J., & Le Maître, E. (2016). Immunomodulatory effects of nicotine on interleukin 1β activated human astrocytes and the role of cyclooxygenase 2 in the underlying mechanism. Journal of Neuroinflammation, 13(1), 256. doi:10.1186/s12974-016-0725-1

Robertson, A. J., Coluccio, A., Jensen, S., Rydlizky, K., & Coffman, J. A. (2013). Sea urchin akt activity is Runx-dependent and required for post-cleavage stage cell division. Biology Open, 2(5), 472–478. doi:10.1242/bio.20133913

Rodriguez, J. J., Terzieva, S., Yeh, C. Y., Gardenal, E., Zallo, F., Verkhratsky, A., & Busquets, X. (2023). Neuroanatomical and morphometric study of S100beta positive astrocytes in the entorhinal cortex during ageing in the 3xTg-Alzehimer’s disease mouse model. Neurosci Lett, 802, 137167. doi:10.1016/j.neulet.2023.137167

Rukstalis, M., Jepson, C., Patterson, F., & Lerman, C. (2005). Increases in hyperactive– impulsive symptoms predict relapse among smokers in nicotine replacement therapy. Journal of Substance Abuse Treatment, 28(4), 297–304. doi:10.1016/j.jsat.2005.02.002

Santello, M., Toni, N., & Volterra, A. (2019). Astrocyte function from information processing to cognition and cognitive impairment. Nat Neurosci, 22(2), 154–166. doi:10.1038/s41593-018-0325-8

Saunders, G. R. B., Wang, X., Chen, F., Jang, S. K., Liu, M., Wang, C., … Vrieze, S. (2022). Genetic diversity fuels gene discovery for tobacco and alcohol use. Nature, 612(7941), 720–724. doi:10.1038/s41586-022-05477-4

Schildge, S., Bohrer, C., Beck, K., & Schachtrup, C. (2013). Isolation and Culture of Mouse Cortical Astrocytes. Journal of Visualized Experiments(71), 50079. doi:10.3791/50079

Sharma, G., & Vijayaraghavan, S. (2001). Nicotinic cholinergic signaling in hippocampal astrocytes involves calcium-induced calcium release from intracellular stores. Proceedings of the National Academy of Sciences of the United States of America, 98(7), 4148–4153. doi:10.1073/pnas.071540198

Shashoua, V. E., Hesse, G. W., & Moore, B. W. (1984). Proteins of the brain extracellular fluid: evidence for release of S-100 protein. J Neurochem, 42(6), 1536–1541. doi:10.1111/j.1471-4159.1984.tb12739.x

Shaw, S., Bencherif, M., & Marrero, M. B. (2002). Janus Kinase 2, an Early Target of α7 Nicotinic Acetylcholine Receptor-mediated Neuroprotection against Aβ-(1–42) Amyloid. Journal of Biological Chemistry, 277(47), 44920–44924. doi:10.1074/jbc.M204610200

Shi, X., Ding, J., Zheng, Y., Wang, J., Sobhani, N., Neeli, P., … Chai, D. (2023). HMGB1/GPC3 dual targeting vaccine induces dendritic cells-mediated CD8+T cell immune response and elicits potential therapeutic effect in hepatocellular carcinoma. iScience, 26(3), 106143. doi:10.1016/j.isci.2023.106143

Shigetomi, E., Bowser, D. N., Sofroniew, M. V., & Khakh, B. S. (2008). Two Forms of Astrocyte Calcium Excitability Have Distinct Effects on NMDA Receptor-Mediated Slow Inward Currents in Pyramidal Neurons. Journal of Neuroscience, 28(26), 6659–6663. doi:10.1523/JNEUROSCI.1717-08.2008

Sigurdsson, S., Alexandersson, K. F., Sulem, P., Feenstra, B., Gudmundsdottir, S., Halldorsson, G. H., … Stefansson, K. (2017). Sequence variants in ARHGAP15, COLQ and FAM155A associate with diverticular disease and diverticulitis. Nat Commun, 8, 15789. doi:10.1038/ncomms15789

Sofroniew, M. V. (2009). Molecular dissection of reactive astrogliosis and glial scar formation. Trends in Neurosciences, 32(12), 638–647. doi:10.1016/j.tins.2009.08.002

Sofroniew, M. V. (2014). Astrogliosis. Cold Spring Harb Perspect Biol, 7(2), a020420. doi:10.1101/cshperspect.a020420

Sole, M., Marazuela, P., Castellote, L., Bonaterra-Pastra, A., Gimenez-Llort, L., & Hernandez-Guillamon, M. (2023). Therapeutic effect of human ApoA-I-Milano variant in aged transgenic mouse model of Alzheimer’s disease. Br J Pharmacol, 180(15), 1999–2017. doi:10.1111/bph.16065

Sun, W., Liu, Z., Jiang, X., Chen, M. B., Dong, H., Liu, J., … Quake, S. R. (2024). Spatial transcriptomics reveal neuron-astrocyte synergy in long-term memory. Nature, 627, 374–381. doi:PMC10937396.

Sweet, L. H., Mulligan, R. C., Finnerty, C. E., Jerskey, B. A., David, S. P., Cohen, R. A., & Niaura, R. S. (2010). Effects of nicotine withdrawal on verbal working memory and associated brain response. Psychiatry Research: Neuroimaging, 183(1), 69–74. doi:10.1016/j.pscychresns.2010.04.014

Tamim, S., Vo, D. T., Uren, P. J., Qiao, M., Bindewald, E., Kasprzak, W. K., … Penalva, L. O. F. (2014). Genomic analyses reveal broad impact of miR-137 on genes associated with malignant transformation and neuronal differentiation in glioblastoma cells. PLoS One, 9(1), e85591. doi:10.1371/journal.pone.0085591

Tarassishin, L., Suh, H.-S., & Lee, S. C. (2014). LPS and IL-1 differentially activate mouse and human astrocytes: Role of CD14: Species-Dependent Astrocyte Immune Activation. Glia, 62(6), 999–1013. doi:10.1002/glia.22657

Taylor, X., Cisternas, P., Jury, N., Martinez, P., Huang, X., You, Y., … Lasagna-Reeves, C. A. (2022). Activated endothelial cells induce a distinct type of astrocytic reactivity. Commun Biol, 5(1), 282. doi:10.1038/s42003-022-03237-8

Teaktong, T., Graham, A., Court, J., Perry, R., Jaros, E., Johnson, M., … Perry, E. (2003). Alzheimer’s disease is associated with a selective increase in alpha7 nicotinic acetylcholine receptor immunoreactivity in astrocytes. Glia, 41(2), 207–211. doi:10.1002/glia.10132

Tsurutani, J., Castillo, S. S., Brognard, J., Granville, C. A., Zhang, C., Gills, J. J., … Dennis, P. A. (2005). Tobacco components stimulate Akt-dependent proliferation and NFkappaB-dependent survival in lung cancer cells. Carcinogenesis, 26(7), 1182–1195. doi:10.1093/carcin/bgi072

Vanrobaeys, Y., Peterson, Z. J., Walsh, E. N., Chatterjee, S., Lin, L. C., Lyons, L. C., … Abel, T. (2023). Spatial transcriptomics reveals unique gene expression changes in different brain regions after sleep deprivation. Nat Commun, 14(1), 7095. doi:10.1038/s41467-023-42751-z

Verkhratsky, A., Parpura, V., Vardjan, N., & Zorec, R. (2019). Physiology of Astroglia. Adv Exp Med Biol, 1175, 45–91. doi:10.1007/978-981-13-9913-8_3

Verkhratsky, A., Rodrigues, J., Pivoriunas, A., Zorec, R., & Semyanov, A. (2019). Astroglial atrophy in Alzheimer’s disease. Pflugers Archiv: European Journal of Physiology, 471(10), 1247–1261. doi:10.1007/s00424-019-02310-2

Verkhratsky, A., Steardo, L., Parpura, V., & Montana, V. (2016). Translational potential of astrocytes in brain disorders. Progress in Neurobiology, 144, 188–205. doi:10.1016/j.pneurobio.2015.09.003

Welsby, P., Rowan, M., & Anwyl, R. (2006). Nicotinic receptor-mediated enhancement of long-term potentiation involves activation of metabotropic glutamate receptors and ryanodine-sensitive calcium stores in the dentate gyrus. European Journal of Neuroscience, 24(11), 3109–3118. doi:10.1111/j.1460-9568.2006.05187.x

West, K. A., Brognard, J., Clark, A. S., Linnoila, I. R., Yang, X., Swain, S. M., … Dennis, P. A. (2003). Rapid Akt activation by nicotine and a tobacco carcinogen modulates the phenotype of normal human airway epithelial cells. The Journal of Clinical Investigation, 111(1), 81–90. doi:10.1172/JCI16147

Wincott, C. M., Abera, S., Vunck, S. A., Tirko, N., Choi, Y., Titcombe, R. F., … Ziff, E. B. (2014). cGMP-dependent protein kinase type II knockout mice exhibit working memory impairments, decreased repetitive behavior, and increased anxiety-like traits. Neurobiology of Learning and Memory, 114, 32–39. doi:10.1016/j.nlm.2014.04.007

Wong, H., Levenga, J., Cain, P., Rothermel, B., Klann, E., & Hoeffer, C. (2015). RCAN1 overexpression promotes age-dependent mitochondrial dysregulation related to neurodegeneration in Alzheimer’s disease. Acta Neuropathol, 130(6), 829–843. doi:10.1007/s00401-015-1499-8

Wong, H., Levenga, J., LaPlante, L., Keller, B., Cooper-Sansone, A., Borski, C., … Hoeffer, C. (2020). Isoform-specific roles for AKT in affective behavior, spatial memory, and extinction related to psychiatric disorders. Elife, 9. doi:10.7554/eLife.56630

Wu, J., Liu, Q., Tang, P., Mikkelsen, J. D., Shen, J., Whiteaker, P., & Yakel, J. L. (2016). Heteromeric alpha7beta2 Nicotinic Acetylcholine Receptors in the Brain. Trends Pharmacol Sci, 37(7), 562–574. doi:10.1016/j.tips.2016.03.005

Xiao, C., Zhou, C. Y., Jiang, J. H., & Yin, C. (2020). Neural circuits and nicotinic acetylcholine receptors mediate the cholinergic regulation of midbrain dopaminergic neurons and nicotine dependence. Acta Pharmacol Sin, 41(1), 1–9. doi:10.1038/s41401-019-0299-4

Xie, Z., Bailey, A., Kuleshov, M. V., Clarke, D. J. B., Evangelista, J. E., Jenkins, S. L., … Ma’ayan, A. (2021). Gene Set Knowledge Discovery with Enrichr. Curr Protoc, 1(3), e90. doi:10.1002/cpz1.90

Xu, S., Ni, H., Chen, H., & Dai, Q. (2019). The interaction between STAT3 and nAChRα1 interferes with nicotine-induced atherosclerosis via Akt/mTOR signaling cascade. Aging, 11(19), 8120–8138. doi:10.18632/aging.102296

Young, J. W., Finlayson, K., Spratt, C., Marston, H. M., Crawford, N., Kelly, J. S., & Sharkey, J. (2004). Nicotine Improves Sustained Attention in Mice: Evidence for Involvement of the α7 Nicotinic Acetylcholine Receptor. Neuropsychopharmacology, 29(5), 891–900. doi:10.1038/sj.npp.1300393

Yu, G., Wang, L., Han, Y., & He, Q. Y. (2012). clusterProfiler: an R package for comparing biological themes among gene clusters. OMICS, 16(5), 284–287. doi:10.1089/omi.2011.0118

Yu, G., Wang, L. G., Han, Y., & He, Q. Y. (2012). clusterProfiler: an R package for comparing biological themes among gene clusters. OMICS, 16(5), 284–287. doi:10.1089/omi.2011.0118

Zeid, D., Kutlu, M. G., & Gould, T. J. (2018). Differential Effects of Nicotine Exposure on the Hippocampus Across Lifespan. Curr Neuropharmacol, 16(4), 388–402. doi:10.2174/1570159X15666170714092436

Zeisel, A., Muñoz-Manchado, A. B., Codeluppi, S., Lönnerberg, P., Manno, G., Juréus, A., … Linnarsson, S. (2015). Brain structure. Cell types in the mouse cortex and hippocampus revealed by single-cell RNA-seq. In.

Zhang, D., Wang, J., Zhou, C., & Xiao, W. (2017). Zebrafish akt2 is essential for survival, growth, bone development, and glucose homeostasis. Mechanisms of Development, 143, 42–52. doi:10.1016/j.mod.2017.01.004

Zhang, J., Han, L., Zhang, A., Wang, Y., Yue, X., You, Y., … Kang, C. (2010). AKT2 expression is associated with glioma malignant progression and required for cell survival and invasion. Oncology Reports, 24(1), 65–72. doi:10.3892/or_00000829

Zhang, K., Förster, R., He, W., Liao, X., Li, J., Yang, C., … Chen, X. (2021). Fear learning induces α7-nicotinic acetylcholine receptor-mediated astrocytic responsiveness that is required for memory persistence. Nature Neuroscience, 24(12), 1686–1698. doi:10.1038/s41593-021-00949-8

Zhang, Z., Ma, Z., Zou, W., Guo, H., Liu, M., Ma, Y., & Zhang, L. (2019). The Appropriate Marker for Astrocytes: Comparing the Distribution and Expression of Three Astrocytic Markers in Different Mouse Cerebral Regions. Biomed Res Int, 2019, 9605265. doi:10.1155/2019/9605265

Zhu, H., Zhuang, D., Lou, Z., Lai, M., Fu, D., Hong, Q., … Zhou, W. (2021). Akt and its phosphorylation in nucleus accumbens mediate heroin-seeking behavior induced by cues in rats. Addiction Biology, 26(5), e13013. doi:10.1111/adb.13013

