## Supplementary material for "AKT2 modulates astrocytic nicotine responses *in vivo*": Table 1

|  | **Figure** | **Homogeneity of variance test** | **Dependent Variable** | **ANOVA / T-Test Results** | **Post hoc testing (Tukey)** | **N/group** |
| --- | --- | --- | --- | --- | --- | --- |
|  | **1A, B** | Bartlett’s P = 0.0787 | Astrocyte Area | F (2, 173) = 31.35  P<0.0001  Power: 1.0000000, Effect Size: 0.5836848 | Saline vs LPS 24 HR: p < 0.0001  Saline vs LPS 72 HR: p < 0.0001  LPS 24 HR vs. LPS 72 HR: p = 0.5348 | 3 animals per group, 20 astrocytes per animal |
|  |  | Spearman’s P < 0.0001 | Total Sholl Intersections | F (2, 4602) = 109.9  P<0.0001  Power: 1.0000000,  Effect Size:  0.1107418 | Saline vs LPS 24 HR: p < 0.0001  Saline vs LPS 72 HR: p < 0.0001  LPS 24 HR vs. LPS 72 HR: p = 0.0606 |  |
| **1C, D** | | Bartlett’s P = 0.9582 | GFAP Sum Intensity | t=0.7450, df=18  P= 0.4659  Power: 0.1085928,  Effect Size:  0.3344010 | N/A | Sal – 9  LPS WT - 11 |

| **Figure** | **Homogeneity of variance test** | **Dependent Variable** | **Anova / T-Test Results** | **Post hoc testing (Tukey** | **N/group** |
| --- | --- | --- | --- | --- | --- |
| **2A, B, C, D** | Bartlett’s P = 0.7476 | Astrocyte Area | Interaction  F (1, 78) = 0.5878, p = 0.4456  Power:  0.9507315  Effect Size:  0.1000000  Treatment F (1, 78) = 20.49, p<0.0001  Power: 0.9999715,  Effect Size: 0.5903925  Genotype  F (1, 78) = 5.192, p =0.0254  Power:  0.8047128,  Effect Size: 0.2971997 | Saline  WT v. A2K: p = .7009  LPS WT v. A2K p = 0.1539  Saline WT vs LPS A2K: p<0.0001  WT  Saline vs LPS: p=0.05  Saline A2K vs. LPS WT: p<0.4018  A2K  Saline vs LPS: p = 0.0016 | 3 animals per group, 20 astrocytes per animal |
|  | Spearman’s P = 0.8100 | Total Sholl Intersections | Interaction  F (27, 780) = 3.759, P<0.0001  Power: 0.8766741  Effect Size: 0.1  Treatment  F (3, 780) = 21.05  P<0.0001  Power: 0.6438766, Effect Size: 0.0769850  Sholl Distance  F (9, 780) = 165.5, p<0.0001  Power: 1.0000000, Effect Size: 0.3954043 | WT Saline vs. WT LPS: p= 0.0489,  WT Saline vs. KO Saline: p = 0.0104  WT Saline vs. KO LPS: p <0.0001  WT LPS vs. KO Saline: p=0.9510  WT LPS vs KO LPS: p<0.0001  KO Saline vs. KO LPS: p=0.0001 |  |
| **2 A** | P <0.0001 | AKT2 Sum Intensity | Welch’s T test, t=10.39, df=8.000, p<0.0001  Power: 1.0000000  Effect Size;  4.8922365 | N/A | WT - 9, *Akt2* cKO - 9 |

| **Figure** | **Homogeneity of variance test** | **Dependent Variable** | **Anova / T-Test Results** | **Post hoc testing (Tukey)** | **N/group** |
| --- | --- | --- | --- | --- | --- |
| **3 A,B,C,D** | Bartlett’s P = 0.0056 | Astrocyte Area | Treatment  F (2, 319) = 17.34, p<0.0001  Power: 0.9998402  Effect Size: 0.3313447 | Saline vs 0.09 mg/kg nicotine: p = 0.0020  Saline vs. 0.2 mg/kg nicotine: p<0.0001  0.09 mg/kg nicotine vs. 0.2 mg/kg nicotine: p=0.0395 | 3 animals per group, 20 astrocytes per animal |
|  | Bartlett’s P<0.0001 | Total Sholl Intersections | Treatment  F (2, 11414) = 147.2, p<0.0001  Power: 1.0000000  Effect Size: 0.0912994 | Saline vs 0.09 mg/kg nicotine: p<0.0001  Saline vs 0.2 mg/kg nicotine: p<0.0001  0.09 mg/kg nicotine vs 0.02 mg/kg nicotine: p<0.0001 |  |
|  | Bartlett’s P = 0.8343 | Cells per mm^2^ | Treatment  F (2, 159) = 0.3757,  P=0.6874  Power: 0.1100969  Effect Size:  0.0683247 | N/A |  |

| **Figure** | **Homogeneity of variance test** | **Dependent Variable** | **Anova / T-Test Results** | **Post hoc testing (Tukey)** | **N/group** |
| --- | --- | --- | --- | --- | --- |
| **Fig 4 A/B** | P = 0.9538 | GFAP Sum Intensity | t=2.152, df=16,  P = 0.0470  Power:  0.5260919  Effect Size:  1.0160753 | N/A | Saline N = 9  Nicotine N = 9 |
|  | Bartlett’s P=0.4252 | GFAP Sum Intensity | Between Groups:  F (2, 33) = 0.6229, p=0.5426  Power: 0.1462353  Effect Size: 0.1870095 | N/A | N=12 for all groups |
| **Fig 4 C** | P = 0.5037 | EAAT2 Sum Intensity | t=7.303, df=8,  P < 0.0001  Power:  0.9999934  Effect Size:  4.7441083 | N/A | Saline N = 4  Nicotine N = 6 |
| **Fig 4 D** | Bartlett’s P = 0.0096 | pAkt2/Akt2 Sum Intensity | Welsh’s t=2.340, df=14.88, p = 0.0337  Power: 0.5750458  Effect Size: 0.9669523 | N/A | Vehicle N=10  Nicotine N=12 |
|  | Bartlett’s P = 0.4510 | Akt2 Sum Intensity | t=2.122, df=20, p = 0.0465  Power: 0.5338064  Effect Size:  0.9198920 | N/A |  |

| **Figure** | **Homogeneity of variance test** | **Dependent Variable** | **Anova / T-Test Results** | **Post hoc testing (Tukey)** | **N/group** |
| --- | --- | --- | --- | --- | --- |
| **5 A,B,C,D** | Bartlett’s P= 0.6007 | Astrocyte Area | Interaction  F (1, 487) = 16.09, p<0.0001  Power: 0.6413702  Effect Size: 0.1000000  Treatment  F (1, 487) = 0.9427, p=0.3321  Power: 0.0782790  Effect Size: 0.0242057  Genotype  F (1, 487) = 2.323, p=.1281  Power: 0.1247779  Effect Size: 0.0380017 | Nicotine  WT vs Akt2 cKO: p=0.1881  Saccharin  WT v Akt2 cKO:  p<0.001  Nicotine WT vs Saccharin Akt2 cKO: p = 0.9698  Nicotine Akt2 cKo vs WT Saccharin: p = 0.3887  Akt2 cKO  Nicotine vs Saccharin: p= 0.0688  WT  Nicotine vs Saccharin: p < 0.001 | 3 animals per group, 20 astrocytes per animal |
|  | Spearman’s P < 0.0001 | Total Sholl Intersections | Interaction  F (75, 25766) = 4.982, p<0.0001  Power: 1.0000000  Effect Size: 0.1000000  Sholl Radius  F (25, 25766) = 2473,  P<0.0001  Power: 1.0000000  Effect Size: 1.2862326  Group  F (3, 25766) = 62.06, p<0.0001  Power: 1.0000000  Effect Size: 0.0705762 | Akt2 cKO  Nicotine vs Saccharin: p<0.0001  Nicotine  Akt2 cKO vs WT: p<0.0001  Nicotine Akt2 cKO vs. Saccharin WT: p=0.2070  Saccharin Akt2 cKO vs Nicotine WT: p=0.4998  Saccharin  Akt2 cKO vs WT: p<0.0001  WT  Nicotine vs Saccharin: p<0.0001 |  |
|  | Bartlett’s P= 0.6007 | Astrocyte Area | Interaction  F (1, 521) = 12.77, p=0.0004  Power: 0.6704618  Effect Size: 0.1000000    Treatment  F (1, 521) = 4.465, p=0.0351  Power: 0.2656182  Effect Size: 0.05912583    Genotype  F (1, 521) = 42.37, p<0.0001  Power: 1.0000000  Effect Size: 0.4412653 | Nicotine  WT vs Akt2 cKO: p<0.0001    Saccharin  WT v Akt2 cKO:  P=0.1755    Nicotine WT vs Saccharin Akt2 cKO: p = 0.0158    Nicotine Akt2 cKo vs WT Saccharin: p <0.0001    Akt2 cKO  Nicotine vs Saccharin: p= 0.0002    WT  Nicotine vs Saccharin: p < 0.7514 |  |

| **Figure** | **Homogeneity of variance test** | **Dependent Variable** | **Anova / T-Test Results** | **Post hoc testing (Tukey)** | **N/group** |
| --- | --- | --- | --- | --- | --- |
| **6 A,B,C,D,E** | Bartlett’s p = 0.0003 | Astrocyte Area | F (3, 76) = 6.739,  p = 0.0004  Power: 0.9840073  Effect Size: 0.5349442 | Vehicle vs. 1uM:  P=0.9579  Vehicle vs 10uM:  p=0.9743  Vehicle vs. 100uM Nicotine: p = 0.0071  1uM Nicotine vs 10uM Nicotine: p = .9998  1uM Nicotine vs 100uM Nicotine: p = 0.0014  10uM vs. 100uM: p = 0.0019 | 3 cultures per group, 10-15 astrocytes per culture |
|  | Bartlett’s p = 0.1084 | Astrocyte Area | F (3, 67) = 3.774,  P = 0.0145  Power: 0.8062979  Effect Size: 0.4064207 | Vehicle vs. 100uM:  p=0.0258  Vehicle vs Vehicle + MLA:  P > 0.9999  Vehicle vs. 100uM Nicotine + MLA: p = 0.9724  100uM Nicotine vs Vehicle + MLA: p = 0.0280  100uM Nicotine vs 100uM Nicotine + MLA: p = 0.1727  Vehicle + MLA vs. 100uM Nicotine + MLA: p = 0.9723 |  |
|  | Spearman’s P = 0.6122 | Astrocyte Area | Interaction  F (1, 33) = 0.2159,  P = 0.6452  Power: 0.0829487  Effect Size: 0.1000000  Treatment  F (1, 33) = 13.87,  P = 0.0007  Power: 0.9994201  Effect Size: 0.8013669  Genotype  F (1, 33) = 4.733,  P = 0.0369  Power: 0.8235212  Effect Size: 0.4681882 | Vehicle  WT vs A2K: p = 0.2558  WT  Vehicle vs 100uM Nicotine: p = 0.0356  WT Vehicle vs A2K 100um Nicotine: p = 0.0017  Vehicle A2K vs 100uM Nicotine WT: p = 0.6678  A2K  Vehicle vs 100uM Nicotine: p = 0.0969  !00uM Nicotine  WT vs A2K: P = 0.6305 |  |
|  | Bartlett’s P = 0.0001 | Astrocyte Area | F (3, 73) = 6.535,  P = 0.0006  Power: 0.9897363  Effect Size: 0.5666762 | Vehicle vs. 100uM:  p=0.0165  Vehicle vs Vehicle + DHBE:  P = 0.5304  Vehicle vs. 100uM Nicotine + DHBE: p = 0.9386  100uM Nicotine vs Vehicle + DHBE: p = 0.0003  100uM Nicotine vs 100uM Nicotine + DHBE: p = 0.0731  Vehicle + DHBE vs. 100uM Nicotine + DHBE: p = 0.2329 |  |

| **Figure** | **Homogeneity of variance test** | **Dependent Variable** | **Anova / T-Test Results** | **Post hoc testing (Tukey)** | **N/group** |
| --- | --- | --- | --- | --- | --- |
| **7 A,B,C,D,E** | P = 0.2047 | Preference Score | t = 2.701, df = 22, P = 0.0130  Power: 0.8231455  Effect Size:  1.0878182 | N/A | WT Nicotine N=11  Akt2 cKO Nicotine N = 13 |
|  | P = 0.5530 | Distance Moved | t=0.8399, df=22, P = 0.4100  Power: 0.2002612  Effect Size: 0.3398418 | N/A |  |
|  | P = 0.6064 | ^Time Spent in Unpreferred Chamber^ | t=0.4453, df=22,  P = 0.6604  Power: 0.1113755  Effect Size:  0.1798356 | N/A |  |

| **Figure** | **Number of uniquely mapped reads (in millions)** | **Normalization method** | **# Genes with adjusted p-value < 0.1** |
| --- | --- | --- | --- |
| **8** | Min: 50.91  1st Qu: 53.62  Median: 58.19 Mean: 57.82  3rd Qu: 60.79 Max: 67.16 | DESeq2 | 35 |
|  | Min: 35.17  1st Qu: 36.99 Median: 39.87 Mean: 39.81  3rd Qu: 42.08 Max: 46.10 | DESeq2 | 7575 |

| **Figure** | **Homogeneity of variance test** | **Dependent Variable** | **Anova / T-Test Results** | **Post hoc testing (Tukey)** | **N/group** |
| --- | --- | --- | --- | --- | --- |
| **Sup. 2 A,B** | P = 0.2143 | Total Sholl Intersections | t=0.4613, df=40,  P = 0.6471  Power: 0.2098082  Effect Size: 0.3638817 | N/A | 3 animals per group, 20 astrocytes per animal |
|  | Spearman’s P < 0.0001 | Total Sholl Intersections | Interaction  F (75, 10608) = 7.649, P<0.0001  Power: 1.0000000  Effect Size: 0.1000000  Sholl Distance  F (25, 10608) = 600.8, P<0.0001  Power: 1.0000000  Effect Size: 0.5117477  Treatment  F (3, 10608) = 133.2, P<0.0001  Power: 1.0000000  Effect Size: 0.0834609 | Saline LPS vs LPS 24 HR: p<0.0001  Saline LPS vs Saline Nicotine: p > 0.9999  Saline LPS vs Nicotine 0.2 mg/kg 24 HR: p,0.0001  LPS 24 HR vs. Saline Nicotine: p<0.0001  LPS 24HR vs Nicotine 0.2 mg/kg 24 HR: p = 0.0778  Saline Nicotine vs Nicotine 0.2 mg/kg 24 HR: p<0.0001 |  |

| **Figure** | **Homogeneity of variance test** | **Dependent Variable** | **Anova / T-Test Results** | **Post hoc testing (Tukey)** | **N/group** |
| --- | --- | --- | --- | --- | --- |
| **Sup. 4 A** | P = 0.3226 | ALDH1L1 Sum Intensity | t=0.1338, df=10,  P = 0.8962  Power: 0.0517554  Effect Size:  0.0786740 | N/A | Saline N=6, Nicotine 24 HR 0.2 mg/kg N=6 |
|  | P = 0.6758 | EAAT2 Sum Intensity | t=0.4243, df=16,  P = 0.6770  Power: 0.0931577  Effect Size: 0.1998172 | N/A | Saline N=9, Nicotine 24 HR 0.2 mg/kg N=9  Saline N=9, Nicotine 24 HR ).2 mg/kg N=9 |
|  | P = 0.5923 | S100β Sum Intensity | t=0.5719, df=16,  P = 0.5753    Power: 0.0842309  Effect Size: 0.2712637 | N/A |  |
| **Sup. 4 B** | P = 0.5402 | VIMENTIN Sum Intensity | t=0.2432, df=10,  P = 0.8128  Power: 0.0555311  Effect Size: 0.1395065 | N/A | Saline N=6, Nicotine 24 HR 0.2 mg/kg N=6 |
|  | P = 0.9968 | OCCLUDIN Sum Intensity | t=0.6216, df=10,  P = 0.5481  Power: 0.0871719  Effect Size: 0.3588961 | N/A |  |

| **Figure** | **Homogeneity of variance test** | **Dependent Variable** | **Anova / T-Test Results** | **Post hoc testing (Tukey)** | **N/group** |
| --- | --- | --- | --- | --- | --- |
| **Sup. 5 A,B,C** | Spearman’s P = 0.2362 | Cells per mm^2^ | Interaction  F (1, 260) = 0.1054, p=0.7457  Power: 0.3719448  Effect Size: 0.10000000    Treatment  F (1, 260) = 1.854, p = 0.1745  Power: 1.0000000  Effect Size: 0.4317696    Genotype  F (1, 260) = 1.421, p = 0.2343  Power: 0.9999995  Effect Size: 0.3671512 | N/A | 3 animals per group, 20 astrocytes per animal |
|  | Bartlett’s P= 0.4203 | Nicotine Consumption (mg/kg/day) | t=0.2614, df=7, p=0.8013  Power: 0.0563316  Effect Size: 0.1807183 | N/A | WT: N=4  Akt2 cKO: N=5 |
|  | Spearman’s P = 0.0202 | Astrocyte Area | Interaction  F (1, 245) = 10.29, p = 0.0015  Vehicle  F (1, 245) = 10.29, p = 0.0015  Genotype  F (1, 245) =  0.09650, p = 0.09650 | Saline WT vs. Saline cKO: p = 0.2093  Saline WT vs. Saccharin WT: p >0.9999  Saline WT vs Saccharin cKO; p = 0.1545  Saline cKO vs Saccharin WT: p = 0.0750  Saline cKO vs Saccahrin cKO: p < 0.0001  Saccharin WT vs. Saccharin cKO: p = 0.0053 | 3 animals per group, 10-20 astrocytes per animal |
