## Supplementary figures and images for "AKT2 modulates astrocytic nicotine responses *in vivo*"

### Supplemental Figure 1

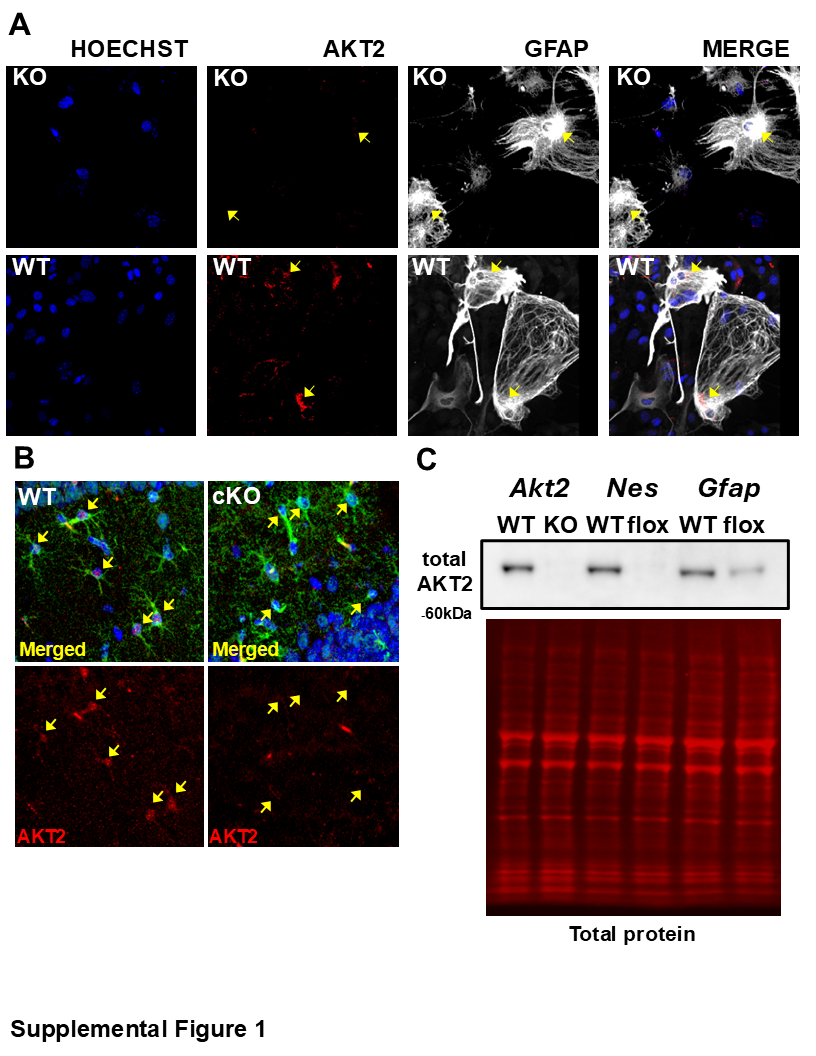

### Supplemental Figure 2

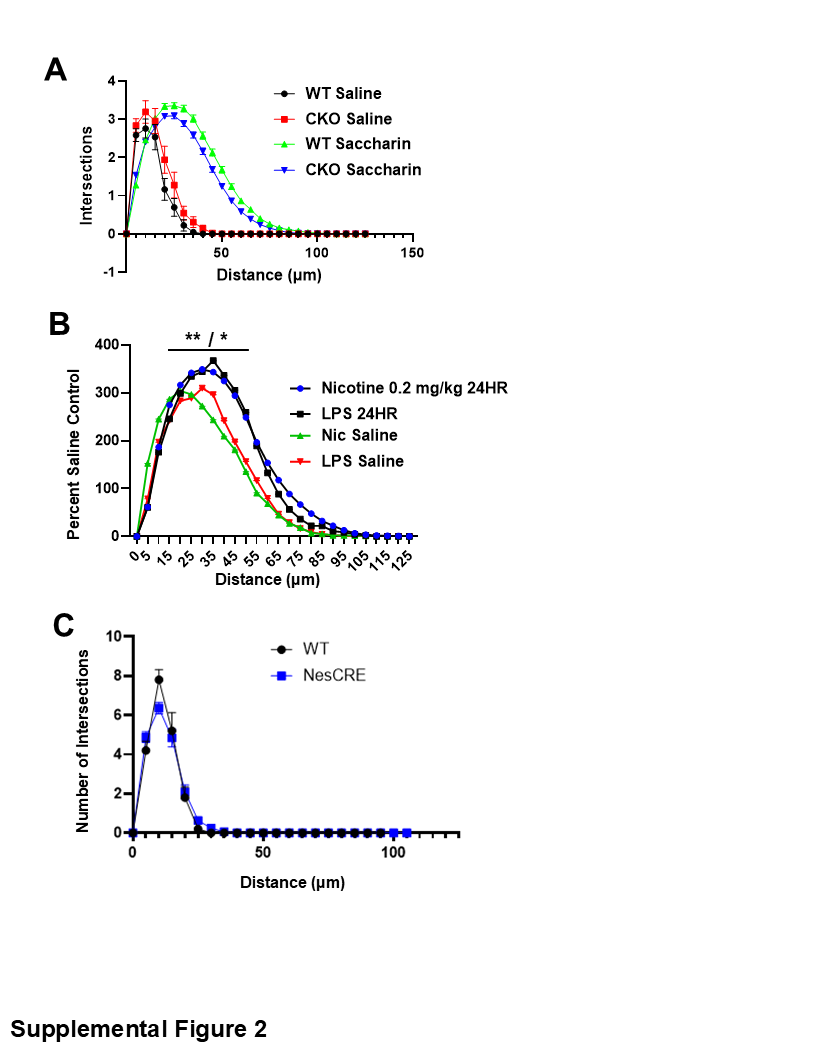

### Supplemental Figure 3

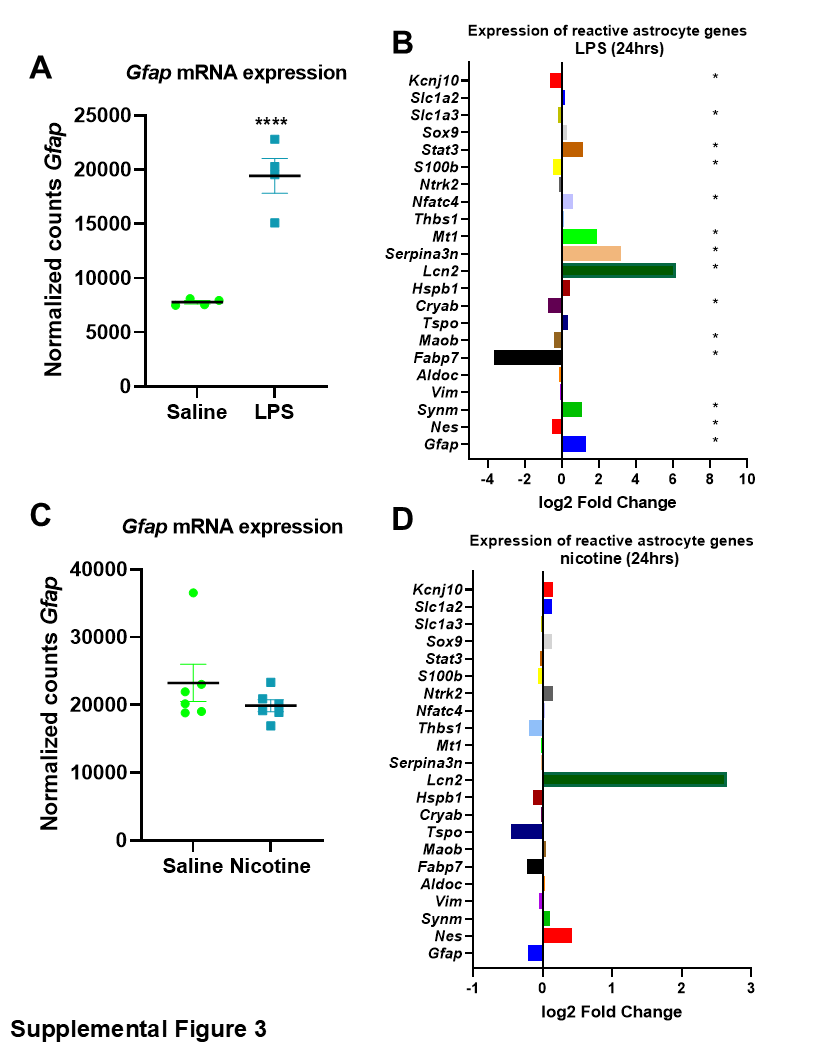

### Supplemental Figure 4

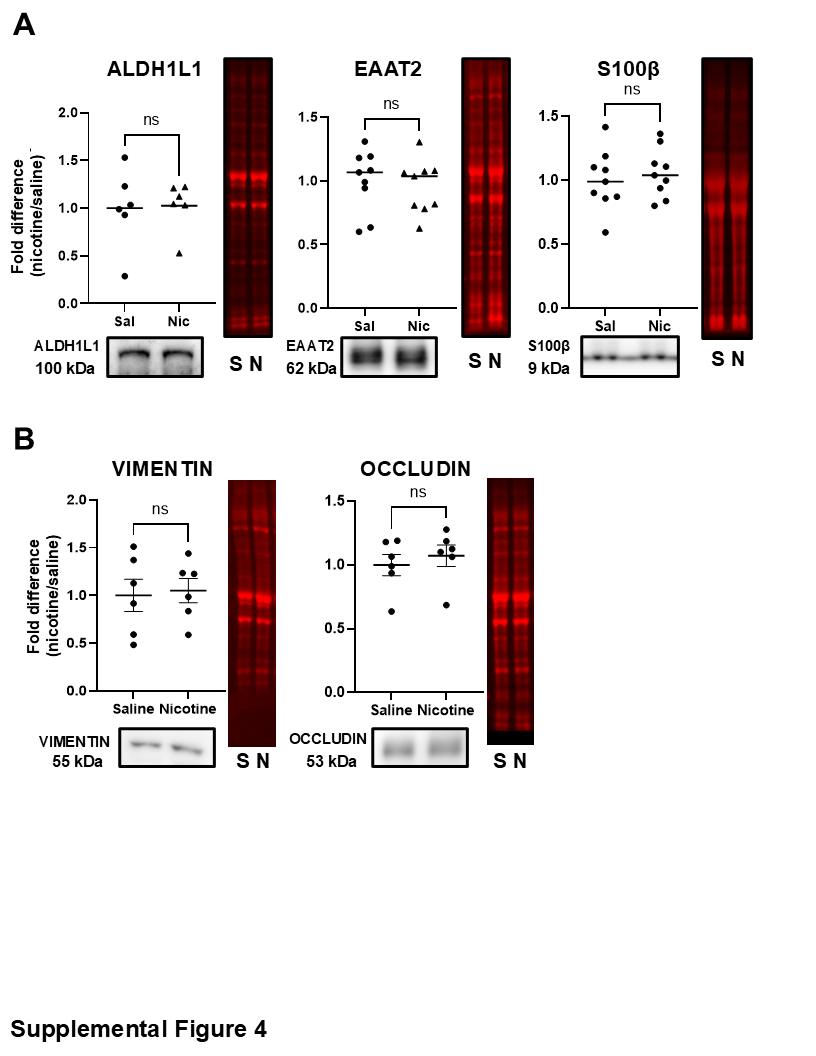

### Supplemental Figure 5

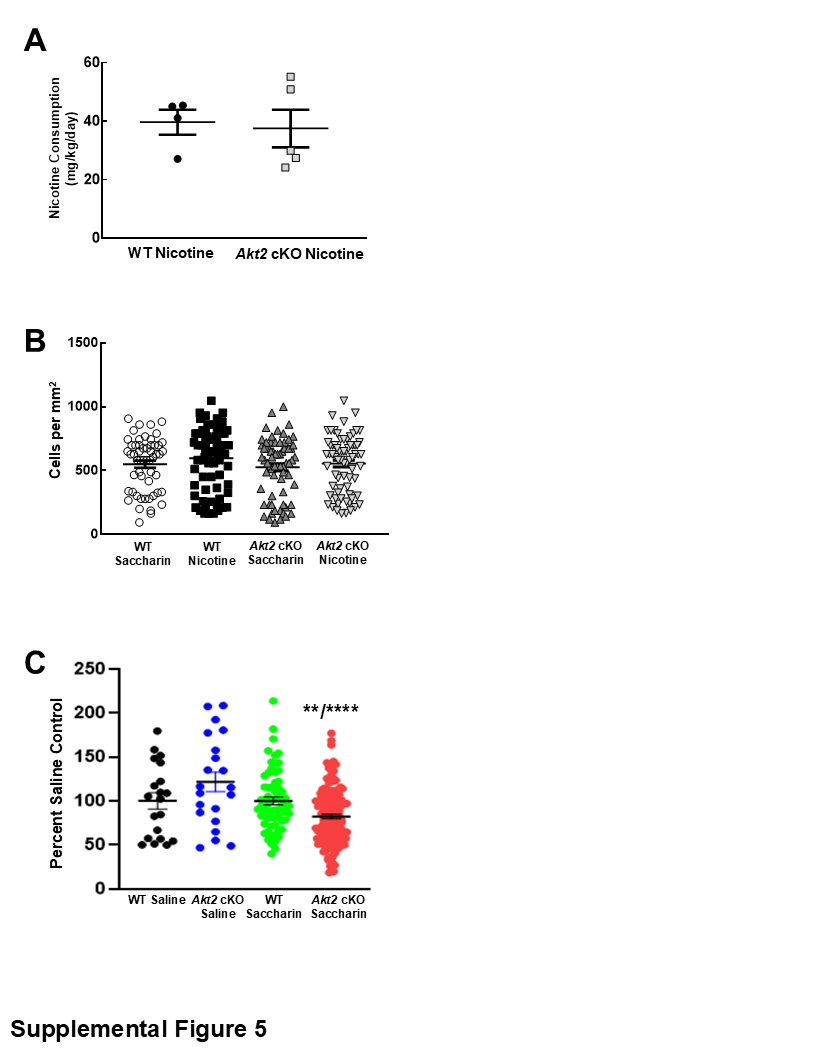
