## Supplemental Figure Legends for "AKT2 modulates astrocytic nicotine responses *in vivo*"

**Supplemental Figure 1.** **Confirmation of *Akt2* deficiency in reagents used for the study.**

**(A**) Staining of *in vitro* primary mouse astrocytes showing loss of AKT2 expression with *Akt2* KO cells compared to WT cells. **(B)** Immunohistochemical staining showing loss of AKT2 expression in hippocampal astrocytes using *Gfap*-*Cre* (G77.6); *Akt2* flox conditional knockout mice. **(C)** Western blot confirmation showing similar loss of AKT2 expression across *Akt2* KO mice (complete null), *Nestin*-*Cre*; *Akt2* flox conditional knockout mice (*Nes*), and *Gfap*-*Cre* (G77.6); *Akt2* flox conditional knockout mice (*Gfap*). Some residual AKT2 protein is visible in *Gfap-Akt2* cKO mice, but verification of AKT2 removal from astrocytes was also performed (B). Protein lysates obtained from whole hippocampal preparations from perfused mice.

**Supplemental Figure 2**. **Imaging and genetic controls for Sholl measurements.**

**(A**) Inter-experimental differences detected between Sholl analysis experiments using different objectives (20X, WT and cKO Saline; 40X WT and cKO Saccharin). **(B)** However, when normalized to saline controls, magnification differences are accounted for. **(C)** No baseline difference in astrocytes is detected with sholl analysis between *Nestin*-*Cre* and WT mice. *p<0.05, **p<.01.

**Supplemental Figure 3**. **GFAP and astrogliosis marker gene expression at 24 h following LPS and nicotine treatments. (A)** Normalized counts for *Gfap* expression 24 hours after *in vivo* LPS exposure, **** padj.<.0001. **(B)** ExpressionLog2 Fold Changes values of astrogliotic markers following LPS exposure, * padj<.01. **(C)** Normalized counts for *Gfap* expression 24 hours after *in vivo* nicotine exposure. **(D)** Log2 Fold Changes values of astrogliotic markers following nicotine exposure**.** *Lcn2* not significant due to low count independent filtering thresholds, otherwise significant.

**Supplemental Figure 4. Western blot analysis of astrocyte activation and blood brain markers following acute i*n vivo* nicotine exposure.** Nicotine-naïve mice were injected i.p. with 0.2 mg/kg nicotine or vehicle (control), and 24 hours later, hippocampal protein lysates were prepared from treated and control mice. Western blot analyses were conducted to compare protein levels of astrocyte activation markers. No differences were detected in **(A)** astrocyte activation markers ALDL1L1, EAAT2, and S100β and **(B)** blood brain markers VIMENTIN and OCCULDIN measured by western blot analysis. N=6 mice (1/hippocampus/mouse) for each treatment group (3M/3F), no sex differences were found, and groups were merged by treatment. Total protein stain next to each quantification, confirm equal loading. T-Test.

**Supplemental Figure 5. Nicotine consumption and cell count controls for chronic nicotine exposure studies. (A)** Drinking consumption between nicotine WT and *Nes*-*Akt2* cKO groups. **(B)** Cell counts for each experimental group. **(C)** Astrocyte area between normalized saline and saccharin conditions. **p < 0.01 Saccharin WT vs Saccharin *Akt2* cKO, ****p<0.0001 Saline *Akt2* cKO vs Saccharin *Akt2* cKO. No significance found for drinking consumption and cell counts, ANOVA, Tukey’s post hoc testing.
